# Hybridization and range expansion in tamarisk beetles (*Diorhabda* spp.) introduced to North America for classical biological control

**DOI:** 10.1101/2021.05.18.444725

**Authors:** Amanda R. Stahlke, Ellyn V. Bitume, A. Zeynep Ozsoy, Dan W. Bean, Anne Veillet, Meaghan I. Clark, Eliza I. Clark, Patrick Moran, Ruth A. Hufbauer, Paul A. Hohenlohe

**Affiliations:** Institute for Bioinformatics and Evolutionary Studies (IBEST), Department of Biological Sciences, University of Idaho, Moscow, Idaho; U.S. Department of Agriculture, Agricultural Research Service (USDA-ARS), Invasive Species and Pollinator Health Research Unit, Albany, CA; U.S. Department of Agriculture, Forest Service (USDA-FS), Pacific Southwest, Institute of Pacific Islands Forestry, Hilo, HI, USA; Department of Biological Sciences, Colorado Mesa University, Grand Junction, CO; Colorado Department of Agriculture, Palisade, CO; Department of Integrative Biology, Michigan State University, East Lansing, MI, USA; Agricultural Biology, Colorado State University, Fort Collins, CO; Graduate Degree Program in Ecology, Colorado State University, Fort Collins, CO

**Keywords:** Biological control, hybridization, range expansion, RADseq, invasion genomics, de novo genome assembly

## Abstract

With the global rise of human-mediated translocations and invasions, it is critical to understand the genomic consequences of hybridization and mechanisms of range expansion. Conventional wisdom is that high genetic drift and loss of genetic diversity due to repeated founder effects will constrain introduced species. However, reduced genetic variation can be countered by behavioral aspects and admixture with other distinct populations. As planned invasions, classical biological control (biocontrol) agents present important opportunities to understand the mechanisms of establishment and spread in a novel environment. The ability of biocontrol agents to spread and adapt, and their effects on local ecosystems, depends on genomic variation and the consequences of admixture in novel environments. Here we use a biocontrol system to examine the genome-wide outcomes of introduction, spread, and hybridization in four cryptic species of a biocontrol agent, the tamarisk beetle (*Diorhabda carinata, D. carinulata, D. elongata*, and *D. sublineata*), introduced from six localities across Eurasia to control the invasive shrub tamarisk (*Tamarix* spp.) in western North America. We assembled a *de novo* draft reference genome and applied RADseq to over 500 individuals from laboratory cultures, the native ranges, and across the introduced range. Despite evidence of a substantial genetic bottleneck among *D. carinulata* in N. America, populations continue to establish and spread, possibly due to aggregation behavior. We found that *D. carinata, D. elongata*, and *D. sublineata* hybridize in the field to varying extents, with *D. carinata* x *D. sublineata* hybrids being the most abundant. Genetic diversity was greater at sites with hybrids, highlighting potential for increased ability to adapt and expand. Our results demonstrate the complex patterns of genomic variation that can result from introduction of multiple ecotypes or species for biocontrol, and the importance of understanding them to predict and manage the effects of biocontrol agents in novel ecosystems.

## 1. Introduction

Human-mediated translocations (e.g., introductions) and climate change have reshaped range limits and previous barriers to gene flow on a global scale (Capinha, Essl, Seebens, Moser, & Pereira, 2015). Conventional wisdom is that high genetic drift and loss of genetic diversity due to repeated founder effects will constrain adaptation, particularly at the expansion front of introduced species or species expanding their ranges due to environmental change (Estoup et al., 2016; Excoffier, Foll, & Petit, 2009; Slatkin & Excoffier, 2012). However, reduced genetic variation can be countered by behavioral aspects of introduced and range-expanding species, such as aggregation, and by admixture with other distinct populations (Estoup *et al*. 2016). Introductions can also provide insights into the stability of cryptic species complexes with allopatric, parapatric, and sympatric native ranges. Introductions can bring such species into contact under novel conditions or secondary contact, representing test cases of ecological speciation (Smith et al., 2018). Furthermore, understanding the genomic consequences of translocations and admixture between introduced populations is key to both preventing the spread of invasive species and to improving the conservation of threatened species (Fauvergue, Vercken, Malausa, & Hufbauer, 2012; McFarlane & Pemberton, 2019; Roderick & Navajas, 2003).

However, the spontaneous nature of accidental introductions makes it difficult to study the outcomes and consequences of eco-evolutionary processes occurring in these systems, since, among other factors, the introduction history and founding population sizes are unknown. Classical biological control programs (hereafter, biocontrol) are essentially planned, intentional invasions. In a typical biocontrol program, highly host-specific natural enemies (agents) are collected in their native range and introduced into a novel environment to control invasive pests (targets) (McFadyen, 1998). Biocontrol systems thus provide an unmatched opportunity to study invasions from a genomic perspective because, compared to their respective target invasive species, biocontrol agents were introduced relatively recently and almost always intentionally, with known source locations, introduction localities, and sometimes known introduction population sizes (Marsico et al., 2010). Despite the opportunity biocontrol agents represent, genomic tools have rarely been used to identify and characterize the consequences of founder effects or evolutionary mechanisms contributing to establishment, persistence, range expansion, or rapid evolution in classical biocontrol agents of invasive species (Hopper et al., 2019; Leung et al., 2020; Muller-Scharer et al., 2020; Sethuraman, Janzen, Weisrock, & Obrycki, 2020; Szűcs, Vercken, Bitume, & Hufbauer, 2019).

Biocontrol scientists have, on several occasions, released several individuals from different locations in the native range, with either known, or inferred, differences in phenotypes. These different populations are referred to as “ecotypes”. Different ecotypes are often released in the hope that some will better match the novel environment (DeBach & Rosen, 1991; Frick, 1970; Room, Harley, Forno, & Sands, 1981; Smith et al., 2018). While this practice may increase the chance of ecological matching across a diverse range of target habitat, it also opens the door for novel phenotypes to arise upon hybridization of different ecotypes. Admixture among different populations, and, at an extreme, hybridization between different species, may present the genetic novelty and diversity necessary to overcome the bottleneck imposed by introduction and adapt, but at the risk of yielding undesirable traits or decreases in fitness (Fauvergue et al., 2012; Kolbe et al., 2004; Lommen, de Jong, & Pannebakker, 2017; Rius & Darling, 2014). The outcomes of multiple agent releases have been understudied, while the consequences of hybridization and admixture among divergent biocontrol agents are even less well understood (but see Szucs, Eigenbrode, Schwarzlander, & Schaffner, 2012; Szűcs et al., 2021; Szűcs, Schaffner, Price, & Schwarzländer, 2012; Szűcs, Schwarzlander, & Gaskin, 2011). Biocontrol efforts could be enhanced if hybridization resulted in increased genetic diversity, providing the raw material for the regional evolution of more efficacious ecotypes, or increased fitness in populations with higher genetic diversity (Bean et al., 2013; Bitume, Bean, Stahlke, & Hufbauer, 2017; Szucs et al., 2012; Szűcs et al., 2021; Szűcs et al., 2012; Tracy & Robbins, 2009). If hybridization resulted in forms with less host specificity than seen in parental forms, for example, biocontrol safety could be compromised; though there have been no documented cases of evolution in fundamental host range (the range of host species on which the agent can complete development) (Van Klinken & Edwards, 2002). Given the potential consequences, post-release monitoring of biocontrol programs involving the release of different populations or species is necessary (Hufbauer, 2008; McFarlane & Pemberton, 2019). Recently developed molecular tools can now also be routinely employed to track biocontrol agents at the population genetics level.

Here, we use a biological control system to understand the consequences of introduction, range expansion, intraspecific admixture, and interspecific hybridization in four introduced closely related species rapidly expanding their ranges. To this end, we produced a *de novo* draft genome assembly of one of the species and used this to (1) identify genetic variation indicating ancestry of the four species, including two ecotype pairs within species, to be able to track their distribution across the introduced range, (2) quantify prevalence and levels of hybridization in the introduced range, and (3) examine the consequences of population bottlenecks and hybridization on genome-wide diversity across broad, landscape-wide expansion fronts and the hybrid zone. We predicted that admixture among populations and hybridizing species would lead to an increase in genetic variation, while isolation of disconnected patches or range expansion would lead to a decrease. We aim here to illustrate how building the molecular genetic foundation to monitor and predict evolution in introduced species can improve our understanding of mechanisms and consequences of range expansion and hybridization in novel environments.

### Study system

The case of *Diorhabda* spp. (Coleoptera: Chrysomelidae), a leaf beetle released to control invasive woody shrubs of the genus *Tamarix* (hereafter, tamarisk), provides a system in which questions relating to the genomic consequences of invasion can be addressed. Tamarisk is native to North Africa and Eurasia and has become invasive in riparian areas across the western United States and northern Mexico. Stands of tamarisk can form dense monotypic thickets that cause substantial economic and environmental damage including increased fire intensity and frequency (Drus, Dudley, Brooks, & Matchett, 2013), increased evapotranspiration (Nagler et al., 2014), diminished soil mycorrhizae critical for native plant species (Meinhardt & Gehring, 2012) as well as a number of other negative impacts on native flora, wildlife habitat and recreation (Di Tomaso, 1998; Gaskin & Schaal, 2002; Shafroth et al., 2005; Zavaleta, 2000). The large extent of the tamarisk invasion, which is estimated to cover at least 360,000 hectares (Nagler, Glenn, Jarnevich, & Shafroth, 2011), coupled with the high value of ecologically-sensitive riparian areas and the cost of conventional control which runs in the millions of dollars (US) per project (Knutson et al., 2019), provided impetus for development and implementation of a biocontrol program.

The first agent released for tamarisk biocontrol was the northern tamarisk beetle, *Diorhabda carinulata*, originally introduced in 2001 from two ecotypes collected in Fukang, China (44.17°N, 87.98°E) and Chilik, Kazakhstan (43.6°N, 78.25°E) (DeLoach et al., 2003). Initial field releases from Fukang and Chilik were performed at eight locations in North America (Fig. 1) (DeLoach et al., 2003) and the species established at northern locations but failed to establish below the 38^th^ parallel in California and Texas (Lewis, DeLoach, Knutson, Tracy, & Robbins et al., 2003) leaving many heavily invaded river systems without a biocontrol option. To address this problem and improve ecological matching across the diverse invaded range (Sands & Harley, 1980), four additional tamarisk-feeding *Diorhabda* populations were collected from ecologically distinct locations in North Africa and Eurasia and introduced primarily in tamarisk-infested areas of western Texas (Knutson et al., 2019; Michels et al., 2013; Tracy & Robbins, 2009). Three ecotypes were elevated to species status in a taxonomic revision based in part on morphology of the genital sclerites (Tracy & Robbins, 2009). As a result, a total of six source populations among four closely related, cryptic species in the genus *Diorhabda* have been released in N. America: *D. carinulata* from Fukang, China and Chilik, Kazakhstan; *D. carinata* from Karshi, Uzbekistan (38.86°N, 65.72°E); *D. sublineata*, from Sfax, Tunisia (34.66°N, 10.67°E); and *D. elongata* from Crete (35.38°N, 24.60°E) and Posidi Beach, Greece (39.96°N, 23.36°E) (Tracy & Robbins, 2009).

**Figure 1.**
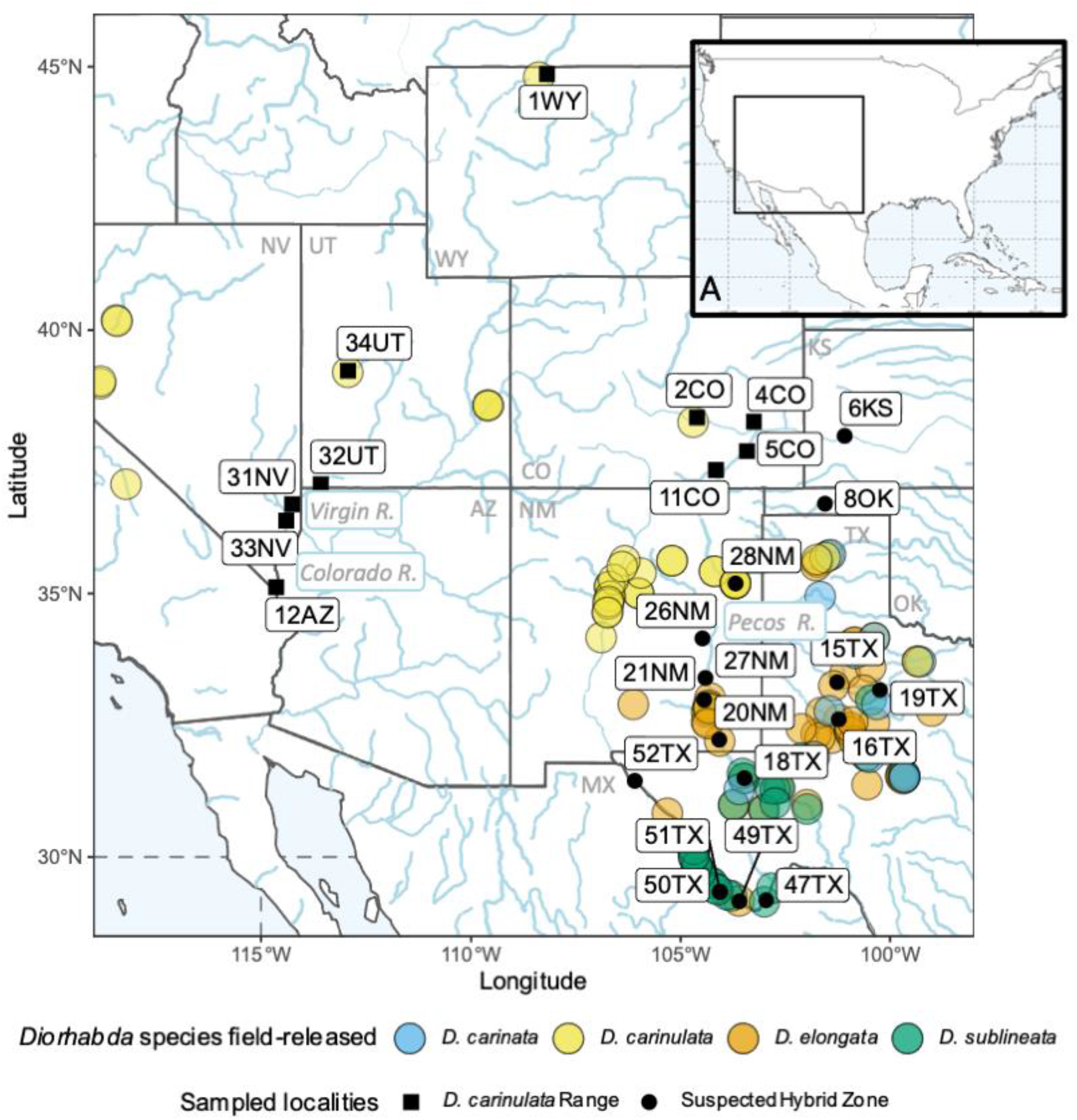
Sampling localities in western North America are indicated in solid black. Squares show both original release sites and localities along the expansion front of *Diorhabda carinulata*. Circles show original release sites and additional locations within the suspected hybrid zone of *D. carinata, D. elongata*, and *D. sublineata*. Transparent filled circles indicate known field-releases (Knutson et al 2019; Supp. Table S3) for each species as indicated in the key. Rivers are drawn in blue. (A) Inset map of N. America centered on the United States and Mexico in top right corner indicates the detailed region presented with a black box.

The *Diorhabda* system is one of the few examples in which contemporary evolution has been demonstrated in a biocontrol agent of invasive plants (see also Szucs et al., 2012; Szűcs et al., 2012; Szűcs et al., 2019 for work on *Longitarsus jacobaeae*). Evolution of response to photoperiod signals enabled rapidly southward expanding *D. carinulata* populations to enter diapause in closer synchrony with the seasonal timing of senescence of tamarisk stands growing in more southern and warmer climates, where the growing season is longer than in the north (Bean, Dalin, & Dudley, 2012; Dalin et al., 2010; Hultine, Bean, Dudley, & Gehring, 2015). Evolution in diapause induction enabled faster range expansion than initially expected (Nagler et al., 2014). Range expansion of *D. carinulata* may not lead to reduced genetic variation at the expansion edge (Slatkin and Excoffier 2012) given negative density-dependent dispersal (Birzu, Matin, Hallatschek, & Korolev, 2019) and migration en masse (‘swarming’) that is common among mobile insects (Sullivan, 1981), especially at range expansion fronts as host resources are depleted and aggregation pheromones draw individuals to mating sites (Cosse, Bartelt, Zilkowski, Bean, & Petroski, 2005).

Additionally, secondary contact between *Diorhabda* species in North America has likely initiated hybridization in this cryptic species complex, uniquely providing a window to the stability of cryptic species upon secondary contact and potentially representing a test case of (non)ecological speciation (Smith et al., 2018). In this case, only *D. carinulata* and *D. carinata* are sympatric in the native range, *D. elongata* is in parapatry with *D. sublineata* to the west and *D. carinata* to the east, and *D. sublineata* and *D. carinata* are the only species completely allopatric (Tracy & Robbins, 2009). While no intermediate forms indicative of hybridization among the four *Diorhabda* species were found in beetles collected from the native range (Tracy & Robbins, 2009), laboratory experiments showed that *D. carinata, D. elongata*, and *D. sublineata* can readily occur and back-cross with viable eggs. In contrast, hybrids and back crosses between *D. carinulata* and the other three species showed significantly reduced egg viability and male sterility (Bean et al., 2013). Later regional studies described intermediate morphotypes between *D. carinata, D. elongata, and D. sublineata* in Texas and surrounding states (Knutson et al., 2019; Michels et al., 2013).

Understanding hybridization, genetic diversity, and range expansion in introduced *Diorhabda* populations is a high priority in management of the tamarisk invasion. A long-standing goal is to evaluate and enhance *Diorhabda* as a tamarisk control option, as the tamarisk invasion and the expense of controlling the shrub at a regional scale has heightened interest in biocontrol among regional resource managers (Bean & Dudley, 2018). Recently, because some native species now utilize tamarisk, including an endangered bird subspecies, the southwest willow flycatcher (SWFL) (*Empidonax traillii extimus*) known to nest in the shrub, a new challenge has been presented to coordinate riparian restoration efforts with declining density of tamarisk brought about by biocontrol with rapidly evolving tamarisk beetles (Hultine et al., 2010; Sogge, Sferra, & Paxton, 2008).

Previous resources to monitor hybridization and range expansion include mitochondrial cytochrome c oxidase I (mt-CO1) haplotypes and morphological markers. These conventional and accessible methods have generally been in agreement for species identification of expanding populations (Bean et al., 2013; Knutson et al., 2019; Ozsoy, Stahlke, Jamison, & Johnson, 2019; Ozsoy, Stahlke, & Johnson, 2018, 2021), but both morphology and CO-1 can lead to incorrect species assignments when hybridization is common, and neither can accurately quantify proportions of ancestry (Rieseberg, Ellstrand, & Arnold, 1993; Wayne & Jenks, 1991) or be used to identify the genetic-basis of ecologically-relevant traits and inform predictions. Thus, molecular genetic analysis at the whole genome level is critical for a more detailed understanding of field populations to inform management strategies.

## 2. Materials and Methods

### 2.1 Whole genome assembly of D. carinulata

We developed a *de novo* draft genome assembly using adults from an inbred line established from field-collected beetles in Lovelock, NV (40.02°N,118.52°W), where *D. carinulata*, originally sourced from Fukang, China, were released in 2001. We sampled reproductively-active males twice from this line, one at the fifth and one at the twelfth generation.

We combined two sequencing approaches for reference genome assembly. First, we extracted genomic DNA from the head, thorax, and dissected testes of the G5 male and constructed a library for whole-genome shotgun sequencing (WGS) using the NEBNext Ultra II DNA library. This WGS library was sequenced in one lane of a MiSeq platform (Illumina) using v3 reagents to produce paired 300-bp reads, resulting in approximately 10.4 million read pairs.

Second, we conducted 10X Chromium sequencing, which produces long-distance synthetically linked reads (Weisenfeld, Kumar, Shah, Church, & Jaffe, 2017). We isolated high molecular weight gDNA from the dissected testes of the G12 male, using a MagAttract kit (Qiagen). The UC Davis Genome Center prepared a Chromium 10X library with v1 chemistry. This 10X library was sequenced on a lane of HiSeq 4000 at UC Berkeley and resulted in 354.48 million read pairs with an average length of 139.50 bp following quality and adapter trimming.

We first assembled the WGS MiSeq 250-bp reads from the G5 male *D. carinulata*. We used a windowed adaptive trimmer, Sickle version 1.33, to remove adapters and low quality reads from this library (Joshi & Fass, 2011) and retained approximately 10.3 million pairs trimmed to an average length of 249.8 bp. We built contigs and scaffolds from these reads with SPAdes version 3.7.1 with one iteration of BayesHammer error correction; k-mer values 21, 33, 55, 77, 99, 127; mismatch careful mode turned on; repeat resolution and mismatchCorrector enabled; and coverage cutoff turned off (Bankevich et al., 2012). Then, we incorporated the 10X Chromium synthetic long reads from the G12 male to scaffold these contigs using the ARCS+LINKS pipeline (Yeo, Coombe, Warren, Chu, & Birol, 2018). Briefly, we extracted barcodes with the 10X software Long Ranger version 2.1.6, aligned these barcoded reads to the SPAdes assembly with BWA-MEM version 0.7.17 (Li, 2013), then supplied the SPAdes assembly and alignments to the ARCS (version 1.0.1) + LINKS (version 1.8.5) pipeline in default mode. ARCS uses the evidence of the synthetic linked reads to construct a graph of linkages for LINKS to then resolve phased scaffolds. We removed contigs of length < 200 bp as required by NCBI. We assessed quality of this assembly in terms of overall contiguity using bbstats.sh from the bbmap suite (Bushnell 2014), as well as the completeness of single-copy conserved orthologous genes using BUSCO version 5.0.0 with the Insecta database (insecta_obd10) composed of 75 species and 1,367 orthologs (Simão, Waterhouse, Ioannidis, Kriventseva, & Zdobnov, 2015).

### 2.2 Sampling and individual genotyping across the native range, introduced range, and lab cultures

Between 2014 and 2017, we field-collected adult beetles to (1) build a reference panel of parental species and ecotypes, (2) characterize their distributions and hybridization in N. America, and (3) examine genomic consequences of range expansion and hybridization. For (1), we collected *D. elongata* and *D. carinulata* in Eurasia near original source collection sites in Greece (Supp. Fig. S1) and China (Supp. Fig. S2). We were not able to collect from the native ranges of *D. sublineata* and *D. carinata*, so we sampled from laboratory colonies derived from the same original collections used for the introductions, maintained at the Palisade Insectary, Colorado Department of Agriculture (see Bean *et al*. 2007 for details of laboratory culturing). To characterize their distribution in N. America (2) and examine the genomic consequences of range expansion and hybridization (3), we collected samples from the sites of the first releases of all four species, along the *D. carinulata* expansion front along the Virgin River, and across the suspected hybrid zone in New Mexico and Texas (Fig. 1, Supp. Table S1, Supp. Figs. S1-S2).

In total we sampled 566 beetles, from 37 locations and two laboratory cultures, for population genomic analysis. At each site, we sampled beetles from trees within a 1 km radius and limited the collection of beetles to no more than 5 individuals per tree where possible. We could not find adults at 19TX and instead collected third-instar larvae which were reared to the adult stage under laboratory conditions. Individuals for 31NV, 32UT, 33NV, and 34UT were sampled from the second generation of laboratory cultures established from these sites. All samples were adult beetles transferred as live individuals to coolers with dry-ice or immediately to a −80°C freezer. All samples were stored at −80°C until DNA extraction.

To prepare restriction-site-associated DNA sequencing (RADseq) libraries, DNA was extracted from individual beetles using a QIAGEN DNeasy Blood and Tissue Kit following the manufacturer’s protocol. The abdomens of all individuals were removed to avoid DNA of developing embryos, gut microbes, or consumed plant material and allow for later morphological characterization. Samples were treated with 4 μL Rnase A (Qiagen) to eliminate RNA contamination. DNA sample concentration was quantified for each individual by fluorometric quantification (Qubit 2.0 HS DNA assay; Invitrogen, Life Technologies, Carlsbad, CA, USA).

In total we prepared 634 individually-barcoded RADseq samples across eight single-digest RADseq libraries using the 8-bp restriction enzyme *SbfI* following the protocol described by Ali et al. (2016). Of those 634, 37 samples were replicated individuals to validate bioinformatic parameter choices and mitigate poor-performing barcodes. Adapter-ligated libraries were multiplexed to achieve approximately 69.7 million reads and paired-end sequenced to 150-bp on an Illumina HiSeq 4000 across two lanes (Vincent J. Coates Genomics Sequencing Laboratory, UC Berkeley). Samples from 47TX-52TX, *D. sublineata* and *D. carinata* cultures, and replicates of ten samples from the first round of sequencing were sequenced in a NovaSeq lane for 80.3 million additional reads. In total, we obtained 149.99 million paired-end reads across all eight RADseq libraries.

We used Stacks 2.5 (Rochette, Rivera-Colón, & Catchen, 2019) to process raw fastq reads, call genotypes, and produce population genetic statistics. First, raw sequencing reads of each library were filtered for PCR duplicates using clone_filter. Then reads were de-multiplexed by individual barcode and re-oriented using the --bestrad flag in process_radtags, allowing for 3 mismatches and discarding reads with low-quality scores (Catchen, Hohenlohe, Bassham, Amores, & Cresko, 2013; Rochette et al., 2019; Stahlke et al., 2020). Each processed sample was then aligned to our *D. carinulata* draft genome using the --very_sensitive flag of bowtie2 version 2.2.9 (Langmead & Salzberg, 2012), and sorted with SAMtools version 1.9 (Li et al., 2009). We called genotypes for all sequenced individuals together in the Stacks 2 module gstacks with the default maruki_low model (Maruki & Lynch, 2017). Then, we required that retained sites were present in the majority of all samples (-R 50) and extracted a random SNP from each ordered locus. Individuals were further filtered to retain those with >4x effective coverage and <75% missing genotypes with VCFtools version 0.1.16 (Danecek et al., 2011). We removed 82 individuals from the dataset with this filter. Finally, genotypes were called again, variant calling format (vcf) files generated, and population genetic summary statistics evaluated using Stacks populations. We refer to the final catalog of SNPs comprising all 552 individuals as the global dataset.

### 2.3 Source Population Ancestry and Hybridization

We used Structure version 2.3.4 (Pritchard, Stephens, & Donnelly, 2000) to identify genetic clusters. Given that founding populations were from distinct sources across Eurasia and the potential for rapid range expansion to lead to dramatic differences in allele frequency among sites, we used the uncorrelated allele frequency model and allowed the alpha parameter to be inferred for each population (Falush, Stephens, & Pritchard, 2003). For each K from 1 to 10, we executed 10 independent runs, allowing a burn-in period of 10,000 steps and 10,000 Markov chain Monte Carlo replicates, and printed the estimation of 90% credible intervals. We used PopHelper 2.3.0 (Francis, 2017) to visualize results and characterize the posterior probability across values of K. After assessing global ancestry assignment with Structure, we checked for relationships between missing genotype rates, alignment rates and individual species ancestry assignment (Supp. Fig. S3).

To identify the genomic characteristics of the four species and track them (1), we compiled published data (Hudgeons et al., 2007; Knutson, DeLoach, Tracy, & Randal, 2012; Knutson et al., 2019; Michels et al., 2013; Pratt et al., 2019) and grey literature describing original releases (directly from the native range) and redistribution efforts (translocations from original release localities) to guide our inference of ancestry assignment (Supp. Table S2). Then we used the Structure ancestry assignments of parental species from lab cultures, single source population release sites (e.g., Fukang ecotype *D. carinulata* at 1WY and Chilik ecotype at 34UT) (Supp. Table S2), and the native range (46CH and 37CR-43GR) to guide our ancestry inference at the remaining localities. Using those diagnostic samples to assign clusters to species identity, we examined the confidence intervals across independent runs to conservatively identify the threshold at which ancestry could be confidently inferred, *q* = 0.067 (i.e. the lower-bound of the 90% credible interval), below which admixture identification could be unreliable and due to technical biases (Caniglia et al., 2020).

Because two of the species, *D. carinulata* and *D. elongata*, were each introduced from two source locations, we examined population substructure within each to identify the distribution of source populations in N. America. We constructed population maps for individuals that had *D. carinulata* or *D. elongata* ancestry (respectively) above the 0.066 threshold, then re-filtered SNPs as above in the Stacks populations module. We then re-ran Structure for each subset of individuals. This secondary analysis also provided a check for sensitivity of population genetic statistics and ancestry assignment to unbalanced sampling in the global Structure analysis (Meirmans, 2019).

Finally, with species identities and population substructure in-hand, we identified hybrids within N. America (2). We classified sampled localities as ‘hybrid’ if the majority of individuals had ancestry from more than one species. We visualized the distributions of q-values (‘hybrid index’) for pairs of inferred ancestral taxa to estimate the degree of back-crossing among pairs (McFarlane & Pemberton, 2019).

Latitudinal variation serves as a proxy for photoperiod, temperature, and host-genotype variation across *Diorhabda* populations. To test for a relationship between ancestry and latitude, we constructed linear models of relevant q-value from latitude for *D. sublineata* ancestry, as the dominant species within the suspected hybrid zone (see Results), and separately, *D. carinulata* source population ancestry to test whether both source populations were represented in southward expansion.

### 2.4 Genome-wide Diversity

To examine the consequences of population bottlenecks and hybridization on genome-wide diversity (3), we quantified genomic differentiation among all individuals within and across sites using population-based measures. We used the populations module of Stacks to calculate *π* (nucleotide diversity), *F*_IS_ (the inbreeding co-efficient), and private alleles. We present *π*and *F*_IS_ calculated from only variant sites, and private alleles were counted among all sites (variant and invariant).

To test whether genetic diversity results (*π, F*_IS,_ and private alleles) were statistically different between hybrids versus pure species and native versus introduced, we performed a one-way ANOVA in R. We grouped samples within localities according to Structure results using the threshold described above. Native range populations included 46CH, 43GR, 44GR, 41 CR, 37CR, 39CR, and 38CR. We excluded laboratory colony populations from this analysis.

To quantify isolation-by-distance (IBD), we calculated the distances between the latitude and longitude coordinates of each sampling locality using the haversine formula and constructed a pairwise distance matrix (Sinnott, 1984). Then, using the vegan package (Oksanen et al., 2019), we conducted a Mantel test with 999 permutations on pair-wise *F*_*ST*_ and distance matrices (Hutchison & Templeton, 1999) for the following groups: (a) introduced *D. carinulata*, (b) native range *D. elongata*, and (c) the suspected hybrid zone. After characterizing IBD, we tested for a linear relationship between latitude of collection site and Chilik ancestry for the *D. carinulata* SNP dataset and, separately, *D. sublineata* ancestry using the global SNP dataset.

To examine the consequences of rapid range expansion in *D. carinulata*, we tested for the signal of asymmetrical range expansion with two origins (1WY and 34UT) across all of introduction sites of *D. carinulata*. Although 1WY is not the source population for *D. carinulata* released in Nevada, Colorado, and Utah, it is the best representative population for Fukang releases (Supp. Table S2). We filtered genotypes in the Stacks populations module for only those populations. We used the rangeExpansion R package version 0.0.0.9000 to estimate the strength of founder effects, the fit of the predicted range expansion dynamic, and the directionality index (ψ), which is a measure of directional clines in allele frequency created by successive colonization events (Peter & Slatkin, 2013).

## 3. Results

### 3.1 Genome assembly

Using SPAdes, we first obtained 21,901 scaffolds with an N50 of 55.67 Kbp, an L50 of 1,790, a total length of 368.964 Mb, and a BUSCO score of 83.9% Complete single-copy orthologs of the 1,367 genes in the BUSCO Insecta set, (86.2% of those represented a single time and 0.7% duplicated), 1.8% fragmented, and 14.3% missing. We improved this assembly by incorporating the 10X Chromium synthetic long reads with the LINKS+ARCS hybrid *de novo* genome assembly approach. This *de novo* draft *D. carinulata* reference genome was composed of 40,962 scaffolds with an N50 of 707.734 Kbp from 108 scaffolds (L50), a total length of 375.976 Mbp, and more complete, single-copy orthologs according to BUSCO with 98.2% complete, (97.4% single, 0.8% duplicated), 1.1% fragmented, and 0.7% missing.

### 3.2 SNP Genotyping

We constructed a total of nine RADseq libraries containing over 552 individuals across lab cultures, the native range, and the introduced range. We removed an average of 39.52% reads identified as PCR duplicates across libraries and retained or recovered a total of 86.28 million reads after initial filtering and demultiplexing. Individual reference alignment rates against the *de novo* draft assembly of *D. carinulata* were 72.77% on average, but we noted two distinct peaks near 75% and 100% that reflected species assignments (Supp. Fig. S3). Effective coverage depth averaged 24.1x after removing individuals with coverage < 4x. In the global SNP dataset, 153,453 loci were merged across paired-end reads for an average locus length of 854.16 bp (std. error = 2.70) and a total of 2,247,632 nucleotide sites. We retained 1,457 SNPs with an average of 8.1% missing data. In the subset of samples with *D. carinulata* ancestry, we retained 2,629 SNPs across 182 samples. In the subset of samples with *D. elongata* ancestry, we retained 1,766 SNPs across 99 samples.

### 3.3 Source Population Ancestry and Hybridization

Structure analysis suggested differential establishment, range expansion, and admixture among all six source populations from the four *Diorhabda* species. We present first the results from the global dataset consisting of all samples from all collections. The greatest change in likelihood, (i.e., the Evanno method) (Evanno, Regnaut, & Goudet, 2005), occurred at *K*=2, splitting *D. carinulata* from the other three introduced *Diorhabda* spp. and hybrids (Supp. Figs. S4-S5). At *K*=4, change in likelihood values plateaued (i.e., the Ln’’(Δ*K*) method; Supp. Fig. S4) and matched species identities for lab cultures, native range, and isolated release sites (Fig. 2). The modal ancestry assignments for *K*=4-8 were identical across replicates after aligning clusters, i.e., no additional clusters were recovered (Supp. Fig. S6). Therefore, we used cluster assignments from *K*=4 to infer parental species and hybridization. Using a threshold of *q*=0.067 for inferring ancestry from a parental species, we found pure ancestry (no interspecific admixture) in 179 *D. carinulata*, 176 *D. sublineata*, 69 *D. carinata*, 93 *D. elongata*. We did not find evidence of hybridization among laboratory cultures or in the source populations for *D. carinulata* (46CH) and *D. elongata* (44GR-41CR).

**Figure 2.**
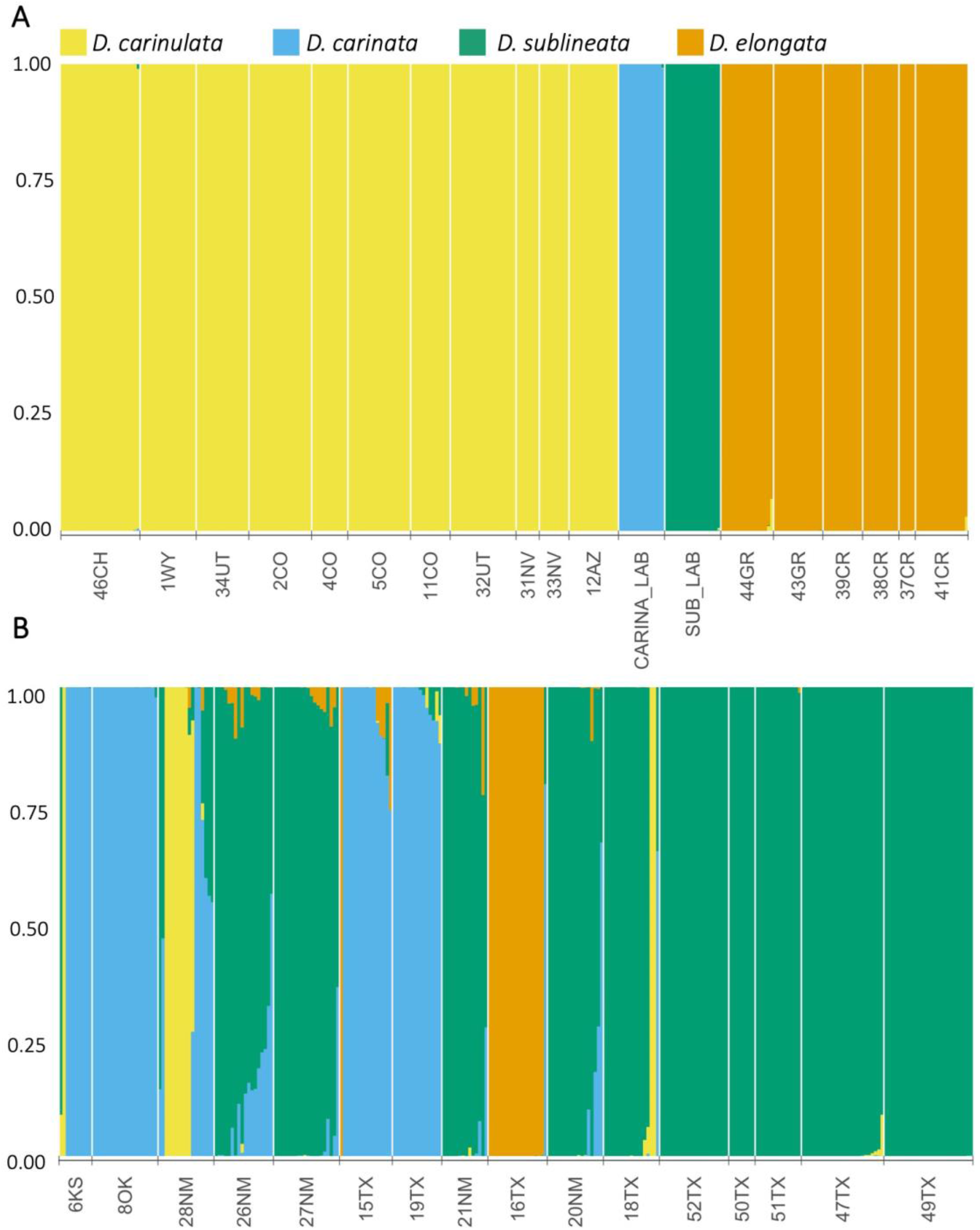
Genetic clustering revealed using Structure (Pritchard et al 2000) for *K*=4 across all species. Each individual sample is represented by a bar. Individuals are grouped by collection site and ordered by localities of Supp. Table 1. Groups from left to right are (A) *D. carinulata* Fukang source collection (46CH, Supp. Figure S2), then north to south from original release sites along expansion front; *D. carinata* and *D. sublineata* lab cultures and native range *D. elongata* collected in Greece (Supp. Figure S1) follow. (B) Individuals collected within the hybrid zone of North America (Figure 1), from north to south.

We further investigated hierarchical substructure among previously identified ecotypes within *D. carinulata* and *D. elongata* (Fig. 3), indicated within the global structure results in a minority of runs (Supp. Fig S5). In both cases, we found that *K*=2 corresponded to the two source populations introduced from each species (Supp. Table S2). We found that the Chilik ecotype, first introduced in 34UT, was the dominant ecotype represented along the Virgin River expansion front (Figs. 1 and 3A). Individuals from sites 5CO and 11CO in eastern Colorado showed admixture with the Chilik ecotype (Fig. 3A), likely due to anthropogenic movement of individuals from western Colorado, which was colonized by individuals from southeast Utah (Supp. Table S2). We detected *D. carinulata* assigned to Fukang ancestry farther east than its suspected range in 6KS, 28NM, and 18TX (Fig. 3A). In *D. elongata*, clusters reflected differentiation between the northern mainland sites and samples collected in Crete (Fig. 3B). We found only the Crete cluster represented in N. America (Fig. 3B). Some individuals we identified as hybrids with other *Diorhabda* species (presented below) showed ancestry from a third cluster (Fig. 3).

**Figure 3.**
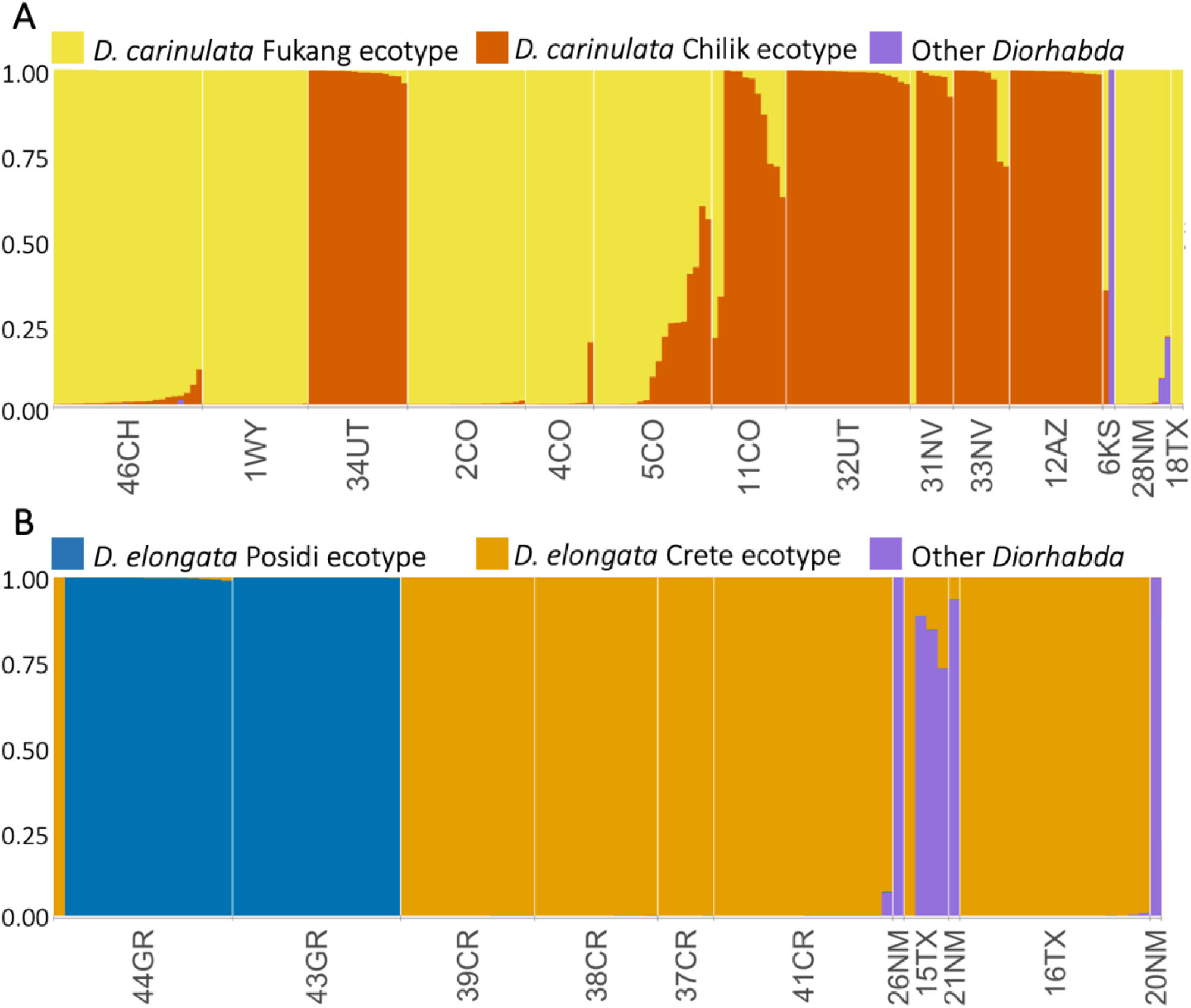
Genetic clustering at K=3 within populations of (A) *D. carinulata* and (B) *D. elongata*.

Most of the hybridization in the global dataset was found at sites near the Pecos River (28NM, 27NM, 26NM, 21NM, 20NM, and 18TX), near where all four species were released (Figs. 1-2, Supp. Table S2) (Knutson et al., 2019). We identified 32 likely hybrids in the suspected hybrid zone: 24 *D. carinata* x *D. sublineata*, three *D. carinata* x *elongata*, three *D. elongata* x *sublineata*, one *D. carinulata* x *D. sublineata*, and one triad hybrid between *D. carinulata* x *D. carinata* x *D. sublineata*. Ranges of q-values for *D. carinata* x *D. sublineata* hybrids included extreme and intermediate values (Supp. Fig. S7A); whereas the other hybrid pairs had ranges less than 0.25 or greater than 0.75 (Supp. Fig. S7B and C). The two putative *D. carinulata* hybrids were from 6KS (G5_rep) and 28NM (G48_rep). G48_rep was assigned largely to *D. sublineata* ancestry with *q*_*sublineata*_ = 0.912, while G5_rep was assigned to tri-specific ancestry with *q*_*carinulata*_ = 0.664, *q*_*sublineata*_ = 0.071, *q*_*carinata*_ = 0.265 (Fig. 2, Supp. Table S3).

We found highly significant linear associations with large residuals between latitude and both *D. sublineata* in the global SNP dataset (Adjusted R^2^ = 0.433; *p* < 0.01) (Supp. Fig. S9A) and the Chilik ecotype in the *D. carinulata* (Adjusted R^2^ = 0.239; *p < 0*.*01*) (Supp. Fig. S9B), demonstrating that latitudinal variation may have influenced differential establishment and spread among genotypes.

### 3.4 Genome-wide diversity

We found significant differences in genetic diversity metrics among hybrid, pure, and native range populations. Nucleotide diversity (*π*) was significantly greater at sites with hybrids (mean = 0.0863) than in either the sites with pure individuals (mean = 0.0335) or from the native range (mean = 0.0381), supporting the prediction that hybridization could increase genetic diversity (Fig. 4). The number of private alleles was greater among sites within the native range (mean = 17.86) than in introduced sites with pure individuals (mean = 7) and hybrids (mean = 7), supporting a bottleneck upon introduction for *D. carinulata* and *D. elongata* (Fig. 4). Sites with hybrids also had a significantly higher average *F*_*IS*_ value (mean = 0.157) than either the sites with pure individuals (mean = 0.0454) or within the native range (mean = 0.027; Fig. 4), consistent with recent hybridization and/or assortative mating among species. All results were significant at *α*= 0.05 (Supp. Table S4).

**Figure 4.**
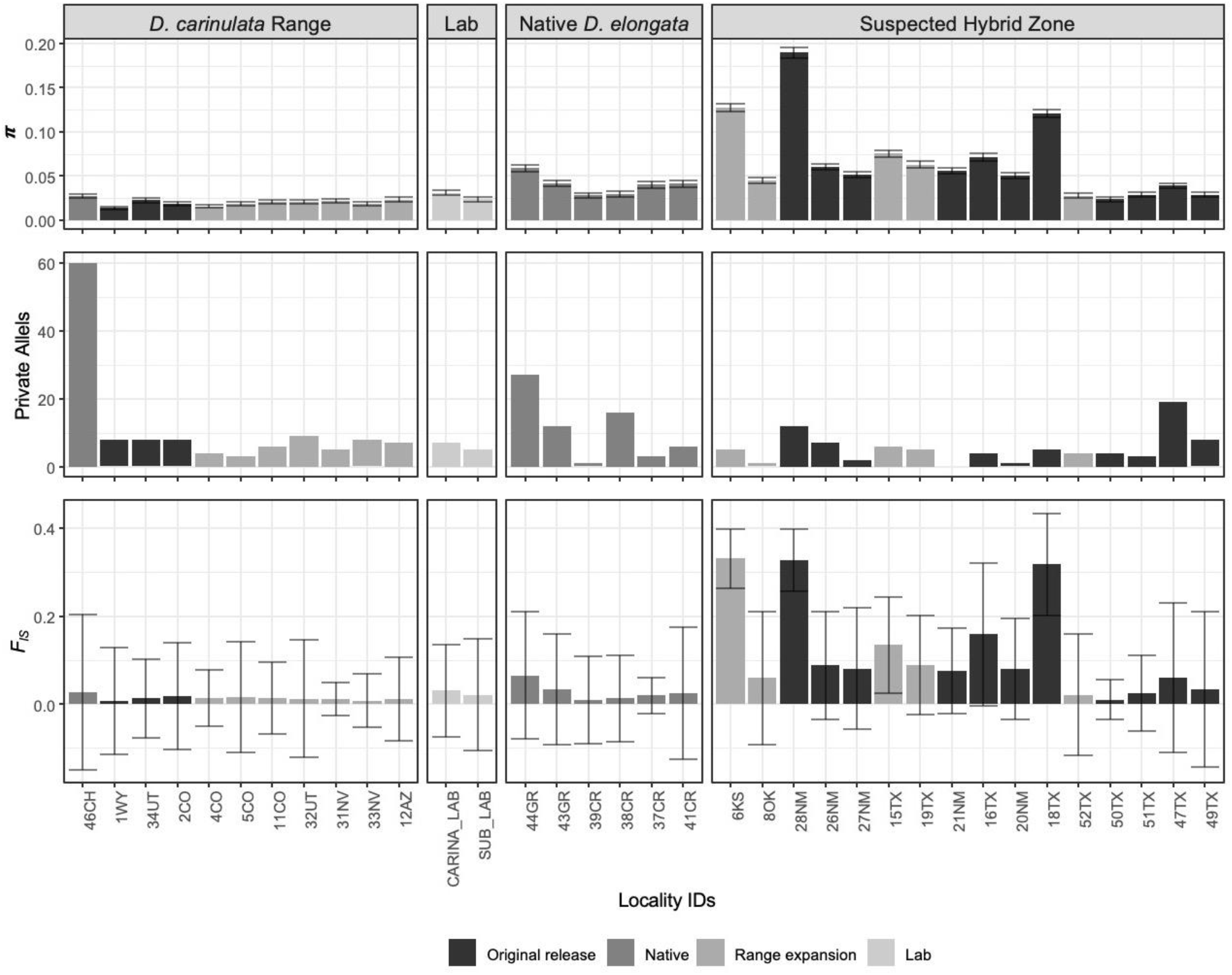
Population genetic statistics including nucleotide diversity ***π*** (top panel), the number of private alleles (middle), and inbreeding coefficient *F*_IS_ (bottom) for the global SNP dataset, consisting of all individuals collected at each locality and grouped (from left to right) by collections within the *D. carinulata* range, laboratory cultures, the native *D. elongata* range, and those within the suspected hybrid zone (ordered as in Figure 2). Shading of individual bars indicates whether the collection was an original release site, site within the native range, natural colonization site, or lab culture (from darkest to lightest). Bars represent +/-the standard error for the respective statistic.

Taking genetic diversity results into a spatial context, we characterized the relationship between divergence and geographic distance among collection sites. Overall evidence for IBD was weak (Fig. 5): Mantel’s r = 0.4481 (*p* = 0.1011) for *D. carinulata*, Mantel’s r = 0.08 (*p* = 0.49167) for native range *D. elongata*, Mantel’s r = 0.5108 (*p* < 0.005) across the suspected hybrid zone. Only the hybrid zone had a positive IBD signal significant at *α*= 0.05. Pairwise comparisons across introduced *D. carinulata* ecotypes and native range *D. elongata* had greater *F*_*ST*_ values and were further apart (Fig. 5, Supp. Fig. S10).

**Figure 5.**
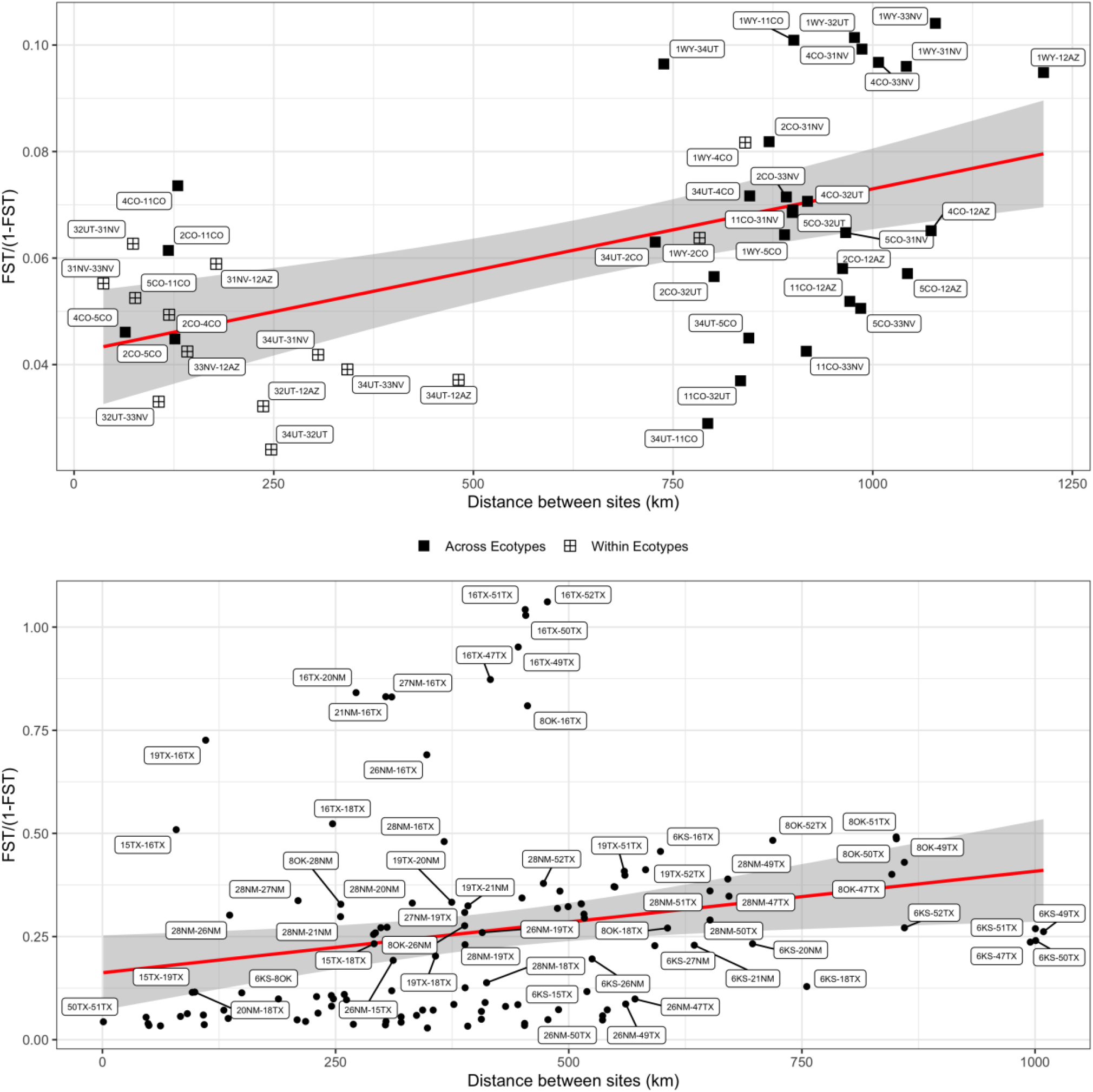
Pairwise *F*_*ST*_ compared to haversine distance between respective sites indicate patterns of isolation-by-distance associated with population structure for (A) *D. carinulata* collected in North America (Mantel’s r = 0.4481, *p* = 0.1011 and (B) across the potential hybrid zone (Mantel’s r = 0.5108, *p* < 0.005). In A, crossed-squares indicate pair-wise comparisons within ecotype, solid squares across ecotype. A linear model for each distribution projected behind points as a red line with standard error in grey.

We detected a weak but significant symmetric range expansion and estimated two distinct origins for *D. carinulata* ecotypes (Supp. Fig. S11; *p* < 0.01). Origin 1 was estimated to be near 1WY (44.86°N, 108.18°W), Origin 2 near the Hoover Dam (35.97°N, 114.64°W), and the origin of their union in central Nebraska (42.63°N, 101.08°W). The directionality index (ψ) was greatest between 32UT (an introduction site of the Chilik ecotype) and 1WY (a Fukang introduction site) at 0.144, supporting two distinct introduced populations, and the least non-negative between 2CO and 18TX (0.0142). Orienting ψ from greatest to least generally reflected suspected expansion fronts within and among the two ecotypes (Supp. Fig. S11).

## 4. Discussion

To better understand the genomic consequences of multi-species introductions and range expansion, we used reference-based population genomics in an exemplar biocontrol system, *Diorhabda* (spp.). This work represents a crucial foundation to advance evolutionary applications by describing the current distribution of species and ecotypes in the introduced range, interspecific hybridization, and impacts on genome-wide diversity. We discuss these results in the context of eco-evolutionary processes, highlight implications for the tamarisk biocontrol program in N. America, and discuss opportunities for further study of this system to improve our understanding of contemporary evolution and biocontrol of invasive plants.

### 4.1 Distribution of parental taxa and hybridization

We found clear evidence for differential establishment of introduced *Diorhabda* populations and spread of these beetles from release sites across the introduced range. Among many original release sites and along suspected colonization routes, individual ancestry assignments were largely composed of the species or population that had been released in the location, suggesting that the extant populations at these release sites remained stable for several generations and naturally expanded along riparian corridors in a predictable pattern. The Fukang release of *D. carinulata* in Lovell, WY (1WY) appeared geographically stable, with uniform ancestry (Fig. 3A) and a lack of inbreeding (Fig. 4) despite being relatively isolated. The Chilik release of *D. carinulata* in Delta, UT (34UT) spread naturally to 12AZ along the Virgin River corridor, and also shows uniform ancestry (Fig. 3A). However, the eco-evolutionary mechanisms driving differential establishment and spread among *Diorhabda* populations will require further study. For example, it is unclear why very little *D. elongata* ancestry was detected in N. America despite many releases in Texas and New Mexico (Fig. 1). The appearance of the *D. carinulata* Fukang ecotype in Kansas (6KS), New Mexico (28NM), and Texas (18TX) (Fig. 3A) was surprising given the lack of establishment reported previously (Bean et al., 2012), although releases of that population did occur near there (Fig. 1). We have preliminary evidence of environmental filtering or adaptation among *Diorhabda* source populations, with genomic clines forming for the *D. carinulata* Chilik ecotype and *D. sublineata* along latitudinal gradients. However, the distributions are confounded by release history (Fig. 1, Supp. Table S2), and the large residuals of this model due to the presence of the *D. carinulata* Fukang ecotype at southern latitudes of NM and TX (Supp. Fig. S9B), suggest that stochastic natural dispersal from release sites or specific genetic variation, rather than broad ecotype identity, could be driving cline formation.

Differential rates of hybridization among *Diorhabda* spp. suggest that there may be several outcomes when multiple closely related taxa are released together, which are likely to depend at least in part upon reproductive barriers. Nonecological speciation theory predicts that when reproductive isolation between taxa is maintained by geography, it is more likely to break down upon secondary contact than when reproductive isolation is maintained by ecological or behavioral differences in sympatric species (Czekanski-Moir & Rundell, 2019). Our data on hybridization frequency supports these predictions. Reproductive isolation between *D. sublineata and D. carinata* was geographic in the native range (Tracy & Robbins, 2009) and has broken down in the introduced range. Reproductive isolation between *D. carinata* and *D. carinulata*, which are sympatric in the native range (Tracy & Robbins, 2009), did not break down upon secondary contact. Although we were not surprised to find hybrids between *D. carinata, D. elongata*, and *D. sublineata* (Bean et al., 2013; Bitume et al., 2017; Knutson et al., 2019), the high abundance (*N*=24) and intermediate distribution of individual q-values among *D. carinata* x *D. sublineata* hybrids in particular suggest a lack of reproductive barriers or even increased fitness relative to parental species, while the distributions of q-values among the other pair-wise hybrids were extreme and could indicate pre- or post-zygotic barriers to hybridization (Supp. Fig. S7)(McFarlane & Pemberton, 2019). In contrast, it is surprising to find *D. carinulata* hybrids (*N*=2) with any of the other species, because laboratory crosses found little to no mating success among those pairs; although *D. carinulata* males could reproduce with females of the other species (Bean et al., 2013). Sex-biased asymmetry of hybridization could be examined by employing existing mt-CO1 efforts with genome-wide ancestry analyses like these (Ozsoy et al., 2019; Ozsoy et al., 2018, 2021; Petit & Excoffier, 2009). We did not recover mtDNA loci in this dataset. The elevated *F*_*IS*_ observed in localities with multiple ancestry (Fig. 4; e.g., 6KS and 28NM with both ‘pure’ samples and hybrids) reflects that parental species were still detected as partially isolated, sympatric populations, not a panmictic population, in the introduced range (i.e. the Wahlund effect) (Waples, 2015). Examining the mechanisms that contributed to speciation of *Diorhabda* in the native range (e.g., divergent environments, ecological interactions, sexual selection) and the role of those barriers in the introduced range present would improve our understanding of the stability of cryptic species broadly and in biocontrol (Fišer, Robinson, & Malard, 2018; Rundle & Nosil, 2005; Smith et al., 2018).

### 4.2 Genomic Diversity during Range Expansion in D. carinulata

The genomic basis of rapid range expansion in *D. carinulata* provides yet another example of the long-held ‘genetic paradox of invasions’, that a reduction of genetic diversity during introduction does not preclude establishment and spread (Estoup et al., 2016). While we found a clear signature of population bottleneck by comparing the number of private alleles between samples from the native range (46CH) to those in N. America (Luikart, Allendorf, Cornuet, & Sherwin, 1998), neither the number of private alleles, nor nucleotide diversity (*π*) declined along range expansion fronts (Fig. 4). The weak signal of asymmetrical range expansion (Supp. Fig. 11) and lack of significant IBD (Fig. 5) suggests that, at least at the time of sampling, population expansion was not unidirectional or smoothly distributed across the landscape, possibly reflecting anthropogenic-mediated dispersal. Alternatively, these results could support a role for negative density-dependent dispersal in preserving genetic diversity (Birzu, Matin, Hallatschek, & Korolev, 2019). If large census and effective population sizes are maintained by this mechanism, it could facilitate rapid local adaptation in novel environments and evolution along expansion fronts by assortative mating for aggregating co-dispersers (Burton, Phillips, & Travis, 2010).

Our range expansion results indicated suggested a contact point for the two introduction sites in Nebraska, far outside the range of geographic possibilities, likely due to the Fukang *D. carinulata* observed in Kansas (6KS) and Texas (18TX) (Fig. 3B). Similarly, several of our sampled sites in the range expansion analysis were not connected by natural dispersal even though they were most genetically similar. To better understand range expansion in this biocontrol system and others like it, a different approach that incorporates anthropogenic movement, spatial distribution of habitat, and direct estimates of dispersal parameters would likely yield a more informative result regarding the genomic mechanisms, routes, and impacts of range expansion. We could build upon this work with biologically-realistic simulations (Haller & Messer, 2019; Landguth et al., 2020) and demographically-informed models using approximate Bayesian computing (Estoup & Guillemaud, 2010) to more accurately assess genomic mechanisms and consequences of rapid range expansion. For example, remote sensing of *D. sublineata* defoliation and expansion has shown that tamarisk continuity and area width predict dispersal distance along a riparian corridor (Ji, Wang, & Knutson, 2017), whereas rangeExpansion assumed a continuous habitat and natural dispersal (Peter & Slatkin, 2013). Further work investigating the ongoing range expansion of *D. carinulata* should also examine possible roles for few loci of large effect (Dlugosch, Anderson, Braasch, Cang, & Gillette, 2015) and plasticity (Bay et al., 2017) operating in the evolution of CDL in *D. carinulata*.

### 4.3 Eco-evolutionary processes influence the impacts of biocontrol

Evolutionary processes have long been of interest in the field of biocontrol (e.g. Simmonds, 1963), and has included efforts to mitigate potential negative effects of losing genetic diversity by augmenting population sizes, and of avoiding ecological mismatch through introducing individuals from deliberately targeted locations in the native range. Despite this long interest, only in recent years are the potential consequences of eco-evolutionary processes on the success of biological control programs being acknowledged and explored (Szűcs et al., 2019). Here we provide further evidence that more detailed and nuanced information, including genomic data, can help us better understand the eco-evolutionary processes occurring among introduced biocontrol agents. Our work specifically documents population expansion despite a dramatic genetic bottleneck, differential establishment and spread among source populations, and differential admixture among those populations.

Widespread hybridization may have implications for both the safety and efficacy of *Tamarix* biocontrol in North America. Because the host plants (biocontrol targets) (*Tamarix chinensis* and *T. ramosissima*) exist primarily in N. America as hybrids (Williams, Friedman, Gaskin, & Norton, 2014), hybridization among *Diorhabda* species may be especially likely to occur and possibly increase fitness (Gilman & Behm, 2011; Seehausen, Takimoto, Roy, & Jokela, 2008). Considering that previous laboratory experiments with hybrids created in the lab showed changes in phenotypes related to both fecundity and host preference (Bitume et al., 2017), the abundance of hybridization we observed warrants further study. These laboratory results have not been verified in the field, and host preference was measured to the third generation (Bitume et al. 2017). A recent regional decline in *Diorhabda* population density, and extirpation from some previously occupied areas, has been noted in Texas, Oklahoma, and Kansas (Knutson et al., 2019). This underscores the possibility that hybrid breakdown could compromise fitness of *Diorhabda* in the field, an interpretation consistent with the observation that up to 57% of field-collected hybrids displayed abnormal genitalic sclerites (Knutson et al., 2019). Testing of these traits (e.g., host choice and fecundity) in field-collected, genotyped populations is critical to better understand changes in risk or efficacy of the biocontrol program due to hybridization. Our dataset could be used to develop a panel of markers for more rapid and cost-effective identification of hybrids with targeted sequencing (e.g., RAD-capture, GTseq) (Meek & Larson, 2019; Reid et al., 2020).

Our population genomics approach presents a much-needed tool to monitor biocontrol releases of multiple populations and cryptic species, highlighted by a notable discrepancy between morphological analyses (Knutson et al., 2019). The laboratory cross found by Bitume et al., (2017) to be the most fecund relative to parental types, *D. carinata* x *D. sublineata*, were the hybrid pairs we found to be most abundant and widely distributed here, but they were not previously described in the morphological analysis of hybridization in this region. One possible explanation is that the sampling design of Knutson et al., (2019) did not include sites farther north into New Mexico, where the bulk of our *D. carinata* x *D. sublineata* hybrids were found. However, we also found some of these hybrids in Texas, in close proximity to the locations sampled by Knutson et al., (2019). These results highlight the value of population genomics to monitor hybridization in cryptic species. Our dataset could be used to improve the accuracy of morphological hybrid identification by validating morphological markers (Griffin, Chandler, Andersen, Havill, & Elkinton, 2020; Padial, Miralles, De la Riva, & Vences, 2010).

Our inferences regarding the mechanisms and impacts of range expansion and hybridization are currently limited because *Diorhabda* collections were sometimes transported without detailed documentation regarding population sizes or population sources. Therefore, we cannot determine based on these data alone whether any hybrid genotype or ancestry combination is more successful without more complete records. For example, we know that many *D. carinulata* releases were made in New Mexico (Supp. Table S2), but the source, precise release sites, precise release numbers, and establishment rates are largely unknown. In general, the practice of anthropogenic movement, often undocumented within management units, presents an interesting tradeoff. On one hand, this increases the availability and likely efficacy of biocontrol across users; but on the other, it makes it more difficult for biocontrol research to understand the patterns of range expansion and adaptation. Evolutionary biologists, biocontrol scientists, and the stakeholders of target invasive species would be well-served with improved records and catalogs of genetic material from biocontrol agent source populations, releases, and follow-up monitoring. Currently there is no entity charged with genomic monitoring of biocontrol releases and these efforts rely on short-term funding and idiosyncratic academic-governmental relationships for each biological system.

The draft assembly of *D. carinulata* is one of only two currently available reference genomes of biocontrol agents of invasive plants, both in the family Chrysomelidae, subfamily Galerucinae (Bouchemousse, Falquet, & Muller-Scharer, 2020), an inviting opportunity for broader comparative work. The resource we developed here is unique in that it can be used for four intentionally released biocontrol agents (the four *Diorhabda* spp.), compared to *Ophraella communa*, which was not intentionally released and is spreading adventively (Müller-Schärer et al., 2014). Nonetheless, only two genomes for any biocontrol agents of invasive plants represents a substantial missed opportunity considering that there are over 332 established invasive plant biocontrol agent species worldwide (Schwarzlander, Hinz, Winston, & Day, 2018). Further development of the *D. carinulata* reference genome, including annotation, would greatly improve our ability to identify SNPs, structural variants, and genes associated with traits related to efficacy and safety as a biocontrol agent; e.g., host choice and diapause induction. Still another opportunity is presented by coupling genomic studies of biocontrol agents with genomic resources from the invasive pests (Lee et al., 2018) to examine co-evolutionary interactions (Sun, Beuchat, & Muller-Scharer, 2020).

The application of genomic approaches in biocontrol systems has the potential to improve both our understanding of contemporary evolutionary processes and management of invasive species and conservation (Leung et al., 2020; Muller-Scharer et al., 2020; Roderick & Navajas, 2003; Sethuraman et al., 2020; Szűcs et al., 2019). Specifically, our results provide a baseline timepoint upon which we can build to further disentangle the mechanisms of rapid range expansion and consequences of hybridization in *Diorhabda*. More generally, we highlight possible predictors of human-mediated translocations and range expansion outcomes, including native-range species boundaries and dispersal behavior, and show how genomic tools in a biocontrol system can test these predictions.

## 5. Data Availability

Code, tabular results, and supplemental tables are hosted at https://github.com/Astahlke/DiorhabaPopulationStructure and will also be posted on Dryad. The reference genome will be available on NCBI under assembly name UIdaho_icDCau_1.0. and is currently being processed by NCBI. Raw sequence reads for the draft reference genome of *D. carinulata* have been deposited to NCBI under PRJNA513507. Raw sequence reads for each RADseq library have been deposited under PRJNA728708 and can be individually demultiplexed with provided barcode files.

## 6. Acknowledgements

The 10X library was constructed at the DNA Technologies and Expression Analysis Cores at the UC Davis Genome Center, supported by NIH Shared Instrumentation Grant 1S10OD010786-01. Sequencing was carried out at the UC Berkeley Vincent J. Coates Genomics Sequencing Laboratory. Genomics and bioinformatics were supported by an Institutional Development Award (IDeA) from the National Institute of General Medical Sciences of the NIH under grant number P30 GM103324. ARS was supported by USDA AFRI NIFA Predoctoral Grant 2020-67034-31888 and the Bioinformatics and Computational Biology Program at the University of Idaho in partnership with IBEST (the Institute for Bioinformatics and Evolutionary Studies). EVB was supported by Postdoctoral fellowship 2015-67012-22931. This research was support by USDA Agriculture and Food Research Initiative grant COLO-2016-09135 to RAH, EB, DWB, and PAH, and RAH acknowledges support from a USDA NIFA Hatch project 1012868. Collections in Xinjiang were supported by the Stillinger Herbarium Expedition Grant and would not have been possible without the help of Daoyuan Zhang and Yan-Feng Chen from the Key Laboratory of Biogeography and Bioresource in Arid Land, Xinjiang Institute of Ecology and Geography, Chinese Academy of Sciences. James Tracy provided coordinates for beetle collections in NM and TX. AZO acknowledges research support from CMU Faculty Professional Development Fund. This work was supported in part by the U.S. Department of Agriculture. USDA is an equal opportunity employer and provider.

