## Supplemental for "Hybridization and range expansion in tamarisk beetles (*Diorhabda* spp.) introduced to North America for classical biological control"

**Tables**

**Supplemental Table S1.** Sampling locality identification codes (Locality ID), site names, latitude and longitude coordinates, dates and sample sizes for each collection effort. Locality IDs include the country or state sampled and symbols for relevant groups as follows: * = native range, Δ = source population, ∇=release sites, ● = hybrid zone, ■ = *D. carinulata* expansion front.

**Supplemental Table S2.** Available data on releases of the different Diorhabda species by year and state. Note: This table does not reflect every release since not all data was accessible at time of publication. Available at https://github.com/Astahlke/DiorhabdaPopulationStructure/blob/master/info/Supp%20tab%203%20Diorhabda%20release%20table.xlsx.

**Supplemental Table S3.** Detailed Structure results for one repetition of $K = 4$ (presented in Figure 2), including sample ID, locality ID, inferred cluster assignments, and 90% credible intervals, available at https://github.com/Astahlke/DiorhabdaPopulationStructure/blob/master/structure_analysis/ancestry_confidenceintervals.csv.

**Supplemental Table S4.** Estimates of $\pi$ (nucleotide diversity), $F_{IS}$ (inbreeding), and number of private alleles for hybrid populations, pure populations, and native populations of *Diorhabda* (spp.) by sample size ($N$). Superscript letters indicate significant differences at $p < 0.05$.

**Figures**

**Supplemental Figure S1.** Sampling sites and original source population locations from the native range of *D. elongata*.

**Supplemental Figure S2.** Sampling sites and original source population locations from the native range of *D. carinulata*.

**Supplemental Figure S3.** Percent of paired reads that mapped concordantly to the *D. carinulata* genome per sample, according to group. A mean of 72.78% is plotted as a horizontal line.

**Supplemental Figure S4.** Change in likelihood values from Structure results for the global SNP dataset, visualized according to four methods (A-D) for values of $K = 1$ to $K = 10$.

**Supplemental Figure S5.** Modal structure results for the global SNP dataset for $K = 2$, $K = 3$, and $K = 4$, from top to bottom.

**Supplemental Figure S6.** Structure results for the global SNP dataset across all ten repetitions of $K = 4$.

**Supplemental Figure S7.** Histograms of individual q-values (as a hybrid index) from modal $K = 4$ Structure results for bi-parental hybrid individuals. Each histogram is symmetric about the 0.50 line and colors indicate the assigned ancestry for the respective cluster. Ordered leading with most abundant pair: A) *D. carinata* x *D. sublineata* hybrids ($n = 24$) have both extreme (near 0 and 1) and intermediate values suggesting a lack of barriers. B) D. elongata x D. carinata hybrids ($n = 3$) and C) *D. elongata* x *D. sublineata* hybrids ($n = 3$) both have more extreme q-values with one likely back-crossed hybrid (0.25, 0.75).

**Supplemental Figure S8.** Change in likelihood values from Structure results for the *D. carinulata* SNP dataset, visualized according to four methods (A-D) for values of $K = 1$ to $K = 10$.

**Supplemental Figure S9.** Change in likelihood values from Structure results for the *D. elongata* SNP dataset, visualized according to four methods (A-D) for values of $K$ = 1 to $K$ = 10.

**Supplemental Figure S10.** Estimated ancestry according to latitude for individuals from sites indicated by color and shape of A) *D. sublineata* from the global set of Structure results and B) *D. carinulata* Fukang ecotype from the *D. carinulata* substructure results. The linear model for each distribution is plotted as a red line with standard error in grey shading.

**Supplemental Figure S11.** Pairwise-$F_{ST}$ compared to haversine distance between respective sites indicate patterns of isolation-by-distance associated with population structure for *D. elongata* collected from the native range in Greece (Mantel's r = 0.086, $p$ = 0.492). Crossed-squares indicate pair-wise comparisons within ecotype, solid squares across ecotype. A linear model for each distribution projected behind points as a red line with standard error in grey to aid in visualization.

**Supplemental Figure S12.** Directionality index ($\psi$) for both *D. carinulata* ecotypes (origins) in North America. Positive stepwise increasing values suggest a directional founding effect. Each ecotype is outlined by a box, with admixture zones (Fig. 3) overlapping.

**Supplemental Table S1.** Sampling locality identification codes (Locality ID), site names, latitude and longitude coordinates, dates and sample sizes for each collection effort. Locality IDs include the country or state sampled and symbols for relevant groups as follows: * = native range, Δ = source population, ∇=release sites, ● = hybrid zone, ■ = *D. carinulata* expansion front.

| Locality ID | Site Name | Latitude (°N) | Longitude (°E) | Date Collected | Sample Size |
|---|---|---|---|---|---|
| ***D. carinulata* sites** | | | | | |
| 46CH*[Δ] | Bitun, China | 47.30 | 87.75 | 7/3/16 | 24 |
| 1WY[∇] | Lovell | 44.86 | -108.18 | 9/3/14 | 17 |
| 34UT[∇] | Delta | 39.23 | -112.93 | 10/15/14 | 16 |
| 2CO[∇] | Fountain Creek | 38.34 | -104.61 | 9/2/14 | 16 |
| 4CO[■] | Adobe Reservoir (Blue Lake) | 38.26 | -103.25 | 9/4/14 | 16 |
| 5CO[■] | SE corner | 37.70 | -103.42 | 8/21/14 | 19 |
| 11CO[■] | Wilkinson | 37.34 | -104.16 | 9/18/14 | 11 |
| 32UT[■] | St. George | 37.07 | -113.58 | 10/15/14 | 10 |
| 31NV[■] | Virgin River, Gold Butte | 36.69 | -114.26 | 10/15/14 | 19 |
| 33NV[■] | Lake Mead, Stewarts Pt | 36.38 | -114.40 | 10/15/14 | 12 |
| 12AZ[■] | Big Bend | 35.12 | -114.64 | 9/18/14 | 20 |
| **Laboratory cultures** | | | | | |
| CARINA_LAB | *D. carinata* Lab Culture | | | 1/7/18 | 20 |
| SUB_LAB | *D. sublineata* Lab Culture | | | 1/7/18 | 7 |
| **Native range *D. elongata*** | | | | | |
| 43GR*[Δ] | Posidi, Greece | 39.97 | 23.37 | 7/6/15 | 9 |

| | | | | | |
|---|---|---|---|---|---|
| 44GR* | Delta Aksiou, Greece | 40.55 | 22.74 | 7/6/15 | 12 |
| 41CR* | Plakias, Crete, Greece | 35.19 | 24.40 | 7/4/15 | 11 |
| 37CR* | Rethimno, Crete, Greece | 35.37 | 24.47 | 7/2/15 | 6 |
| 39CR* | Panaramnos, Crete, Greece | 35.42 | 24.68 | 7/4/15 | 16 |
| 38CR*Δ | Sfkaki, Crete, GR | 35.42 | 24.69 | 7/4/15 | 17 |
| **Suspected Hybrid Zone** | | | | | |
| 6KS• | W Finney County | 37.99 | -101.08 | 8/8/14 | 15 |
| 8OK• | Guymon | 36.70 | -101.55 | 9/16/14 | 18 |
| 28NM• | Tucumcari Lake | 35.19 | -103.69 | 10/6/14 | 20 |
| 26NM• | Lake Sumner | 34.15 | -104.48 | 9/29/14 | 16 |
| 27NM• | Roswell E | 33.40 | -104.41 | 9/29/14 | 15 |
| 15TX• | NE Post | 33.32 | -101.26 | 9/26/14 | 15 |
| 19TX• | Aspermont | 33.17 | -100.24 | 9/27/14 | 18 |
| 21NM•∇ | Artesia Wildlife Reserve | 32.98 | -104.44 | 10/7/14 | 17 |
| 16TX•∇ | Lake JB Thomas | 32.61 | -101.22 | 9/26/14 | 17 |
| 20NM• | Malaga | 32.22 | -104.08 | 9/27/14 | 21 |
| 18TX• | Orla | 31.49 | -103.48 | 9/27/14 | 8 |
| 52TX• | Tornillo | 31.44 | -106.09 | 8/3/17 | 14 |
| 50TX• | Presidio Hwy | 29.34 | -104.07 | 8/3/17 | 25 |
| 51TX• | CO Canyon Boat Ramp | 29.34 | -104.06 | 8/3/17 | 27 |
| 47TX• | Rio Grande Village | 29.18 | -102.96 | 8/3/17 | 14 |
| 49TX∇• | Santa Elana | 29.16 | -103.60 | 8/3/17 | 17 |

**Supplemental Table 3.** Estimates of $\boldsymbol{\pi}$ (nucleotide diversity), $F_{IS}$ (inbreeding), and number of private alleles for hybrid populations, pure populations, and native populations of *Diorhabda* (spp.) by sample size (*N*). Superscript letters indicate significant differences at $p<0.05$.

| | *N* | Mean (95% CI) | *P* |
|---|---|---|---|
| $\boldsymbol{\pi}$ | | | |
| Hybrid | 7 | 0.0863 (0.063, 0.11)[a] | 0.002 |
| Pure | 21 | 0.0335 (0.015, 0.062)[b] | |
| Native | 7 | 0.0381 (0.014, 0.062)[b] | |
| $\boldsymbol{F_{IS}}$ | | | |
| Hybrid | 7 | 0.157 (0.098, 0.217)[a] | 0.004 |
| Pure | 21 | 0.0454 (0.011, 0.080)[b] | |
| Native | 7 | 0.027 (-0.033, 0.0868)[b] | |
| **Private alleles** | | | |
| Hybrid | 7 | 4.71 (-2.63, 12.1)[a] | 0.019 |
| Pure | 21 | 6.24 (2.00, 10.5)[a] | |
| Native | 7 | 17.86 (10.52, 25.2)[b] | |

**Supplemental Figure S1.** Sampling sites and original source population locations from the native range of *D. elongata*.

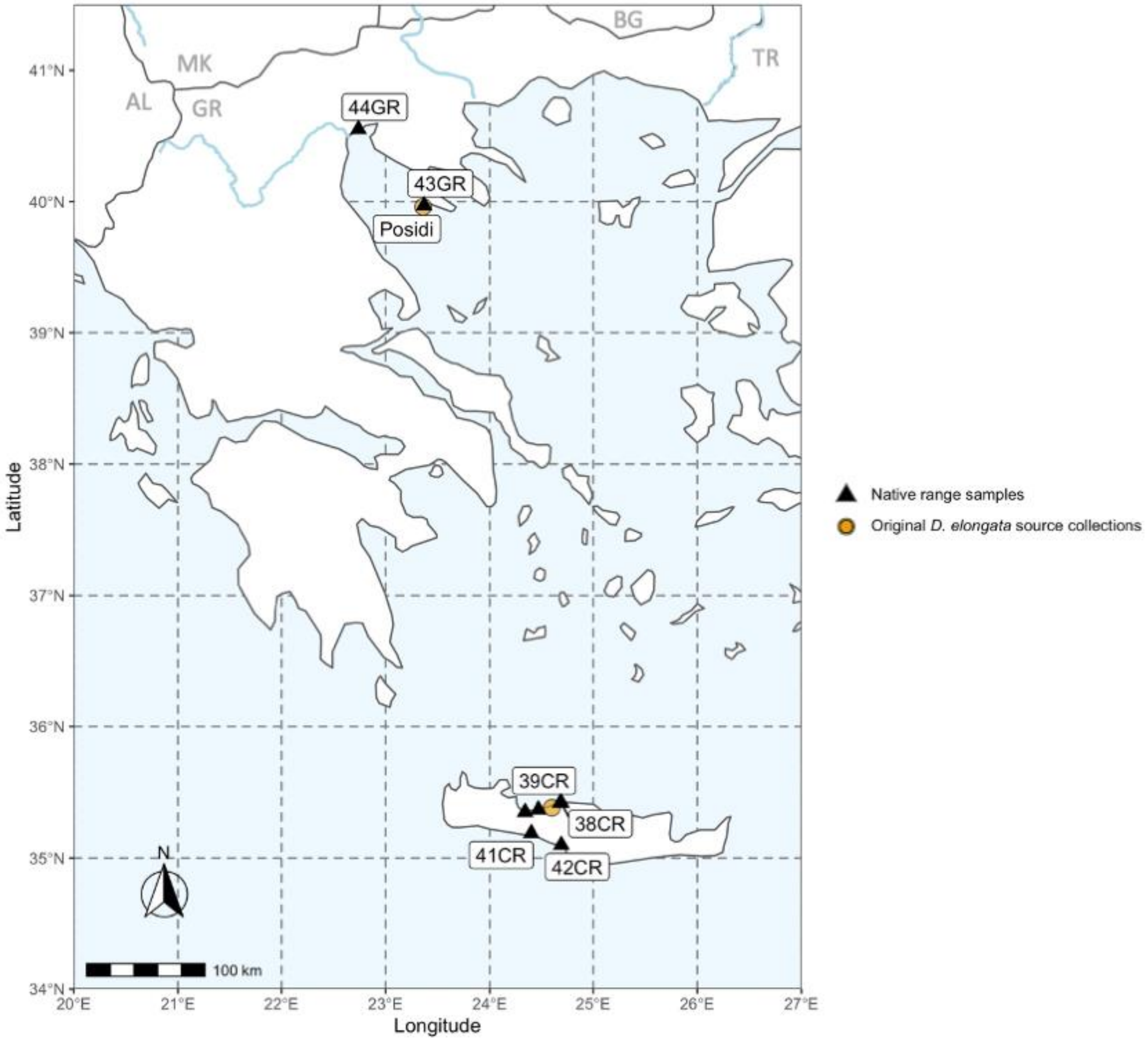

**Supplemental Figure S2.** Sampling sites and original source population locations from the native range of *D. carinulata*.

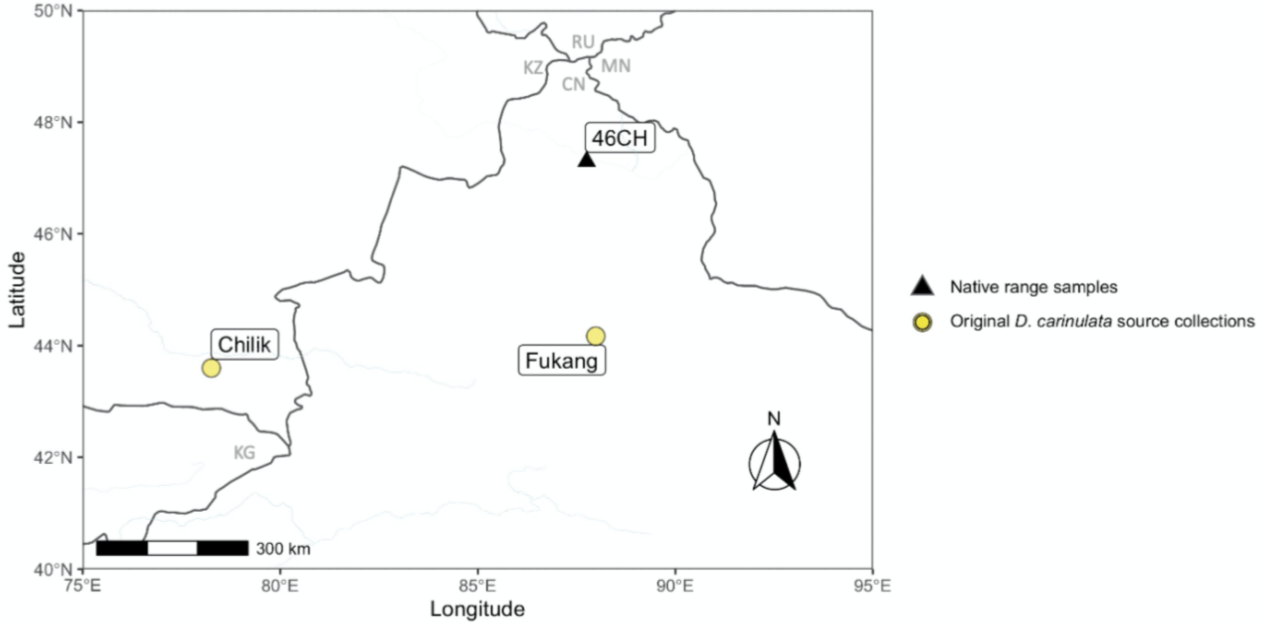

**Supplemental Figure S3.** Percent of paired reads that mapped concordantly to the *D. carinulata* genome per sample, according to group. The mean of 72.78% is plotted as a horizontal line.

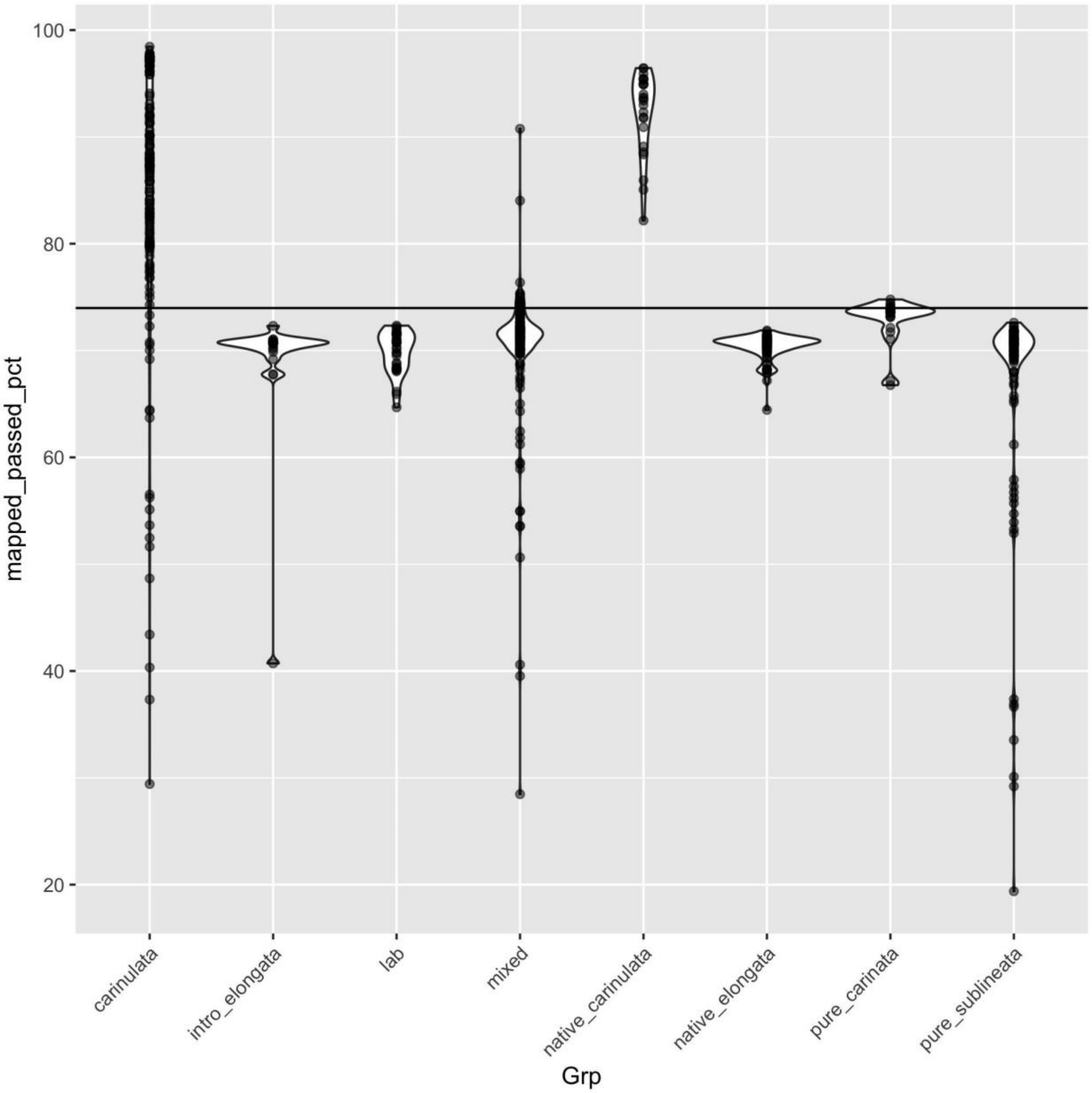

**Supplemental Figure S4.** Change in likelihood values from Structure results for the global SNP dataset, visualized according to four methods (A-D) for values of $K$ = 1 to $K$ = 10.

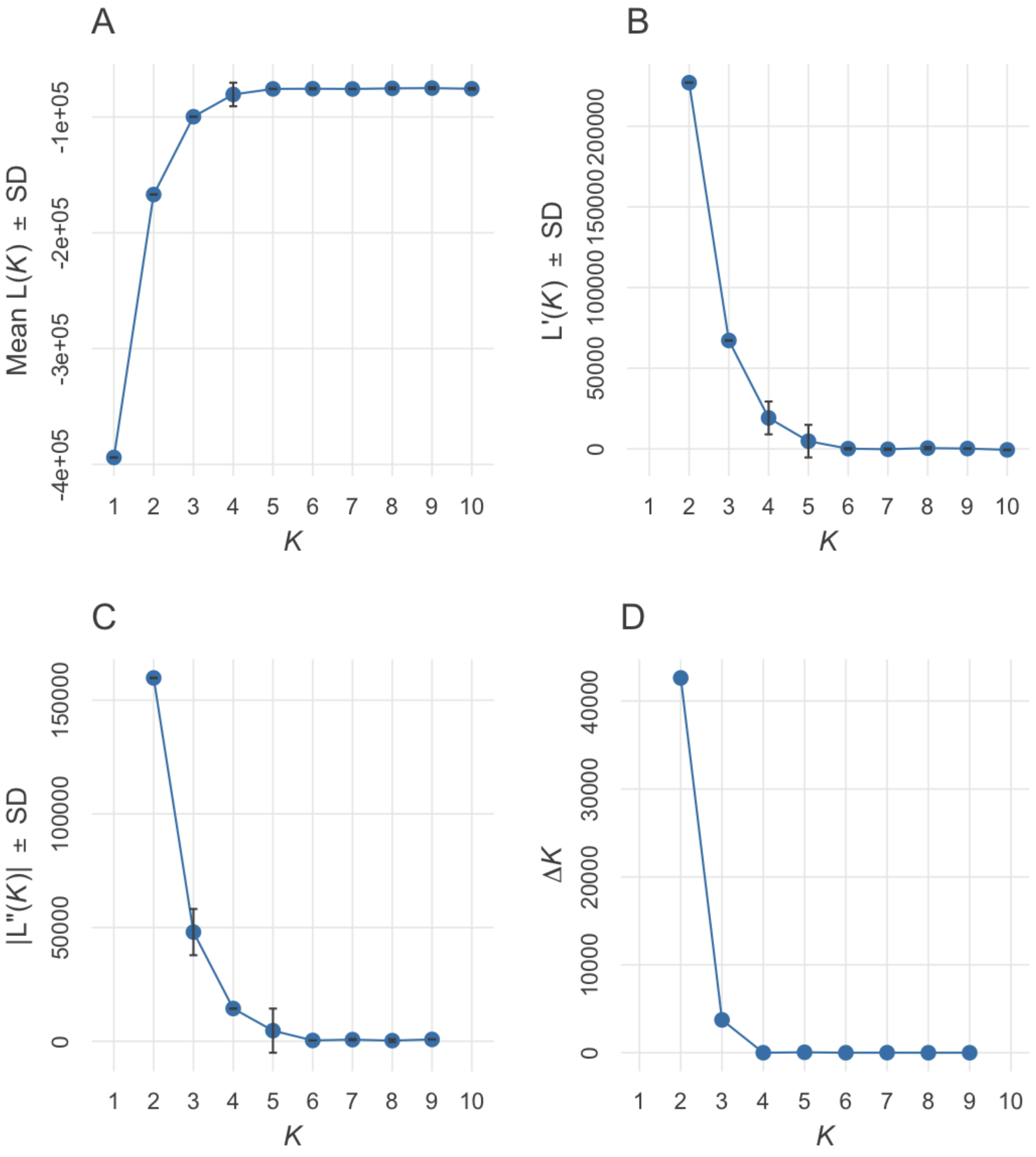

**Supplemental Figure S5.** Modal structure results for the global SNP dataset for K=2, K=3, and K=4, from top to bottom.

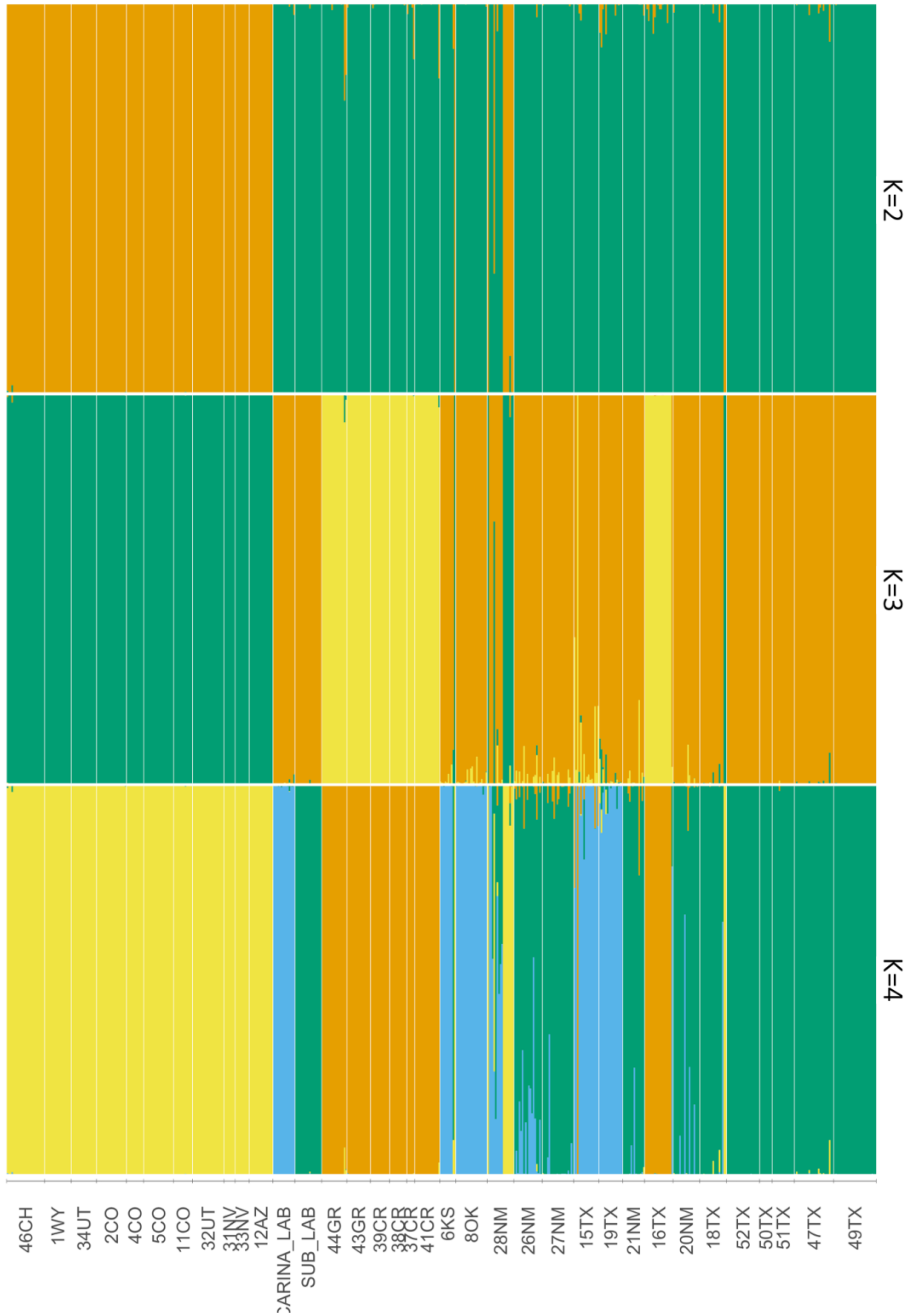

**Supplemental Figure S6.** Structure results for the global SNP dataset across all ten repetitions of K=4.

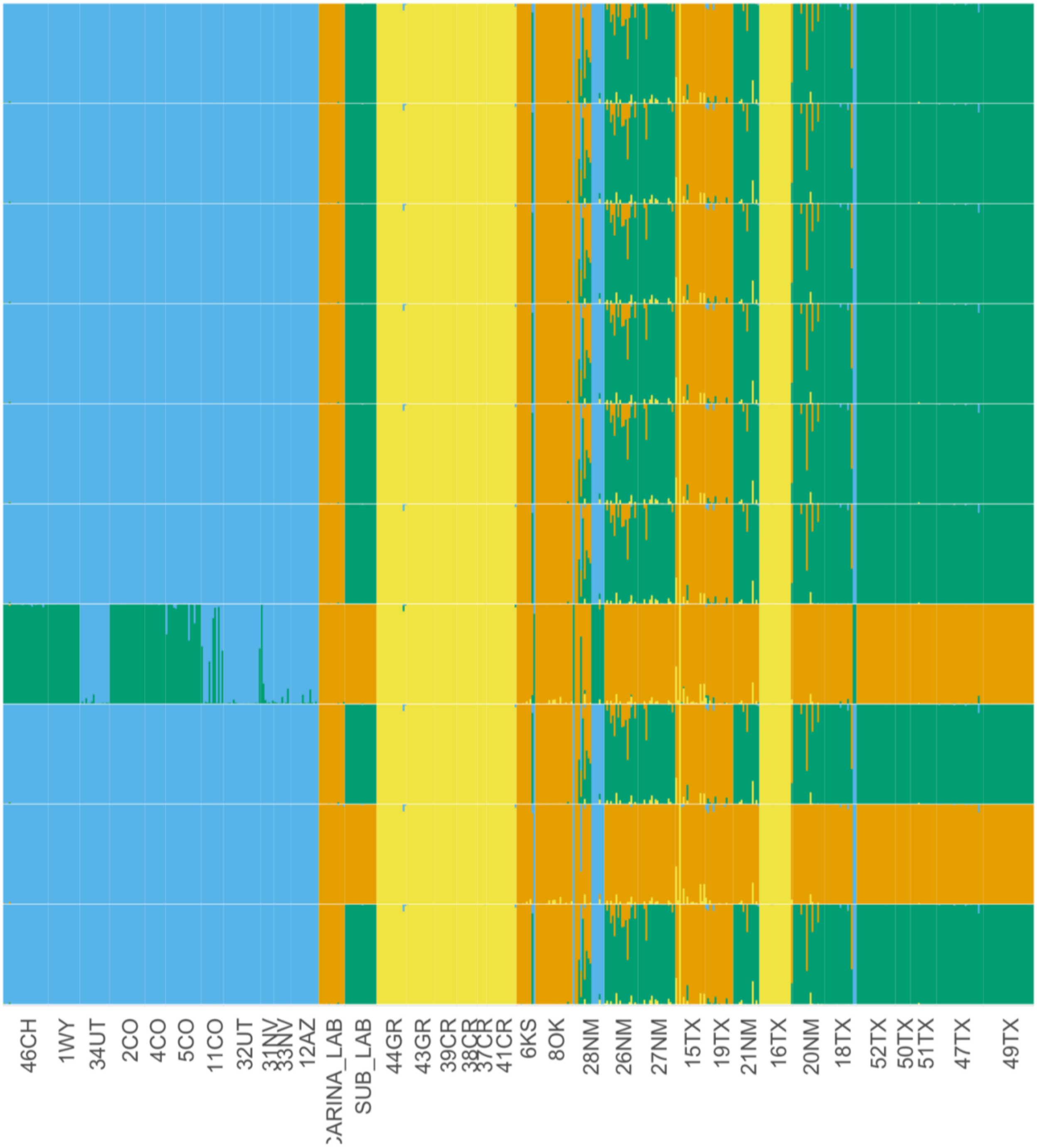

**Supplemental Figure S7.** Histograms of individual q-values (as a hybrid index) from modal *K*=4 Structure (Pritchard et al 2000) results for bi-parental hybrid individuals. Each histogram is symmetric about the 0.50 line and colors indicate the assigned ancestry for the respective cluster. Ordered leading with most abundant pair: A) *D. carinata* x *D. sublineata* hybrids (*n*=24) have both extreme (near 0 and 1) and intermediate values suggesting a lack of barriers. B) *D. elongata* x *D. carinata* hybrids (*n*=3) and C) *D. elongata* x *D. sublineata* hybrids (*n*=3) both have more extreme q-values with one likely back-crossed hybrid (0.25, 0.75).

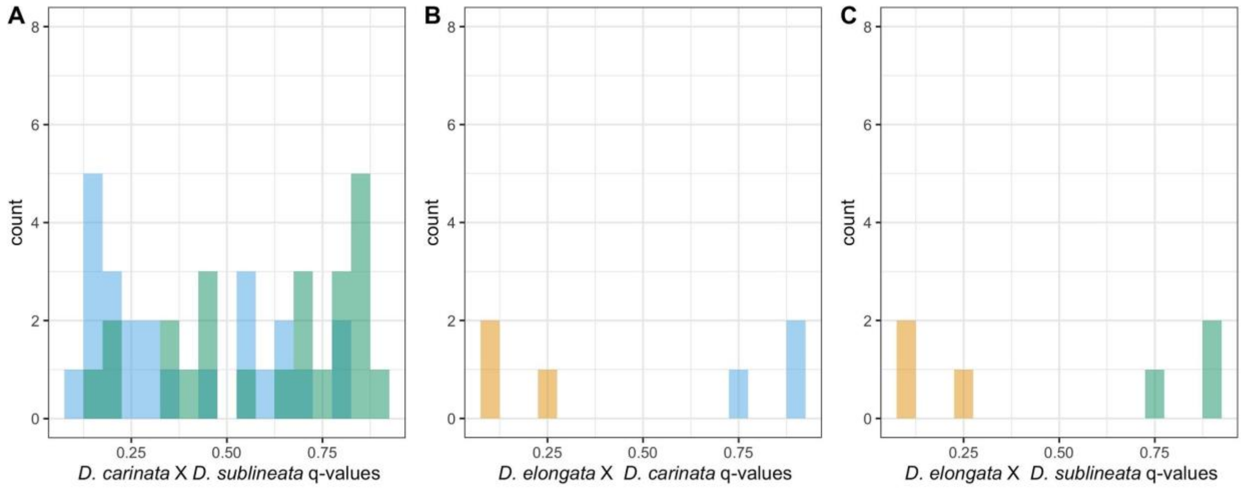

**Supplemental Figure S8.** Change in likelihood values from Structure results for the *D. carinulata* SNP dataset, visualized according to four methods (A-D) for values of $K$ = 1 to $K$ = 10.

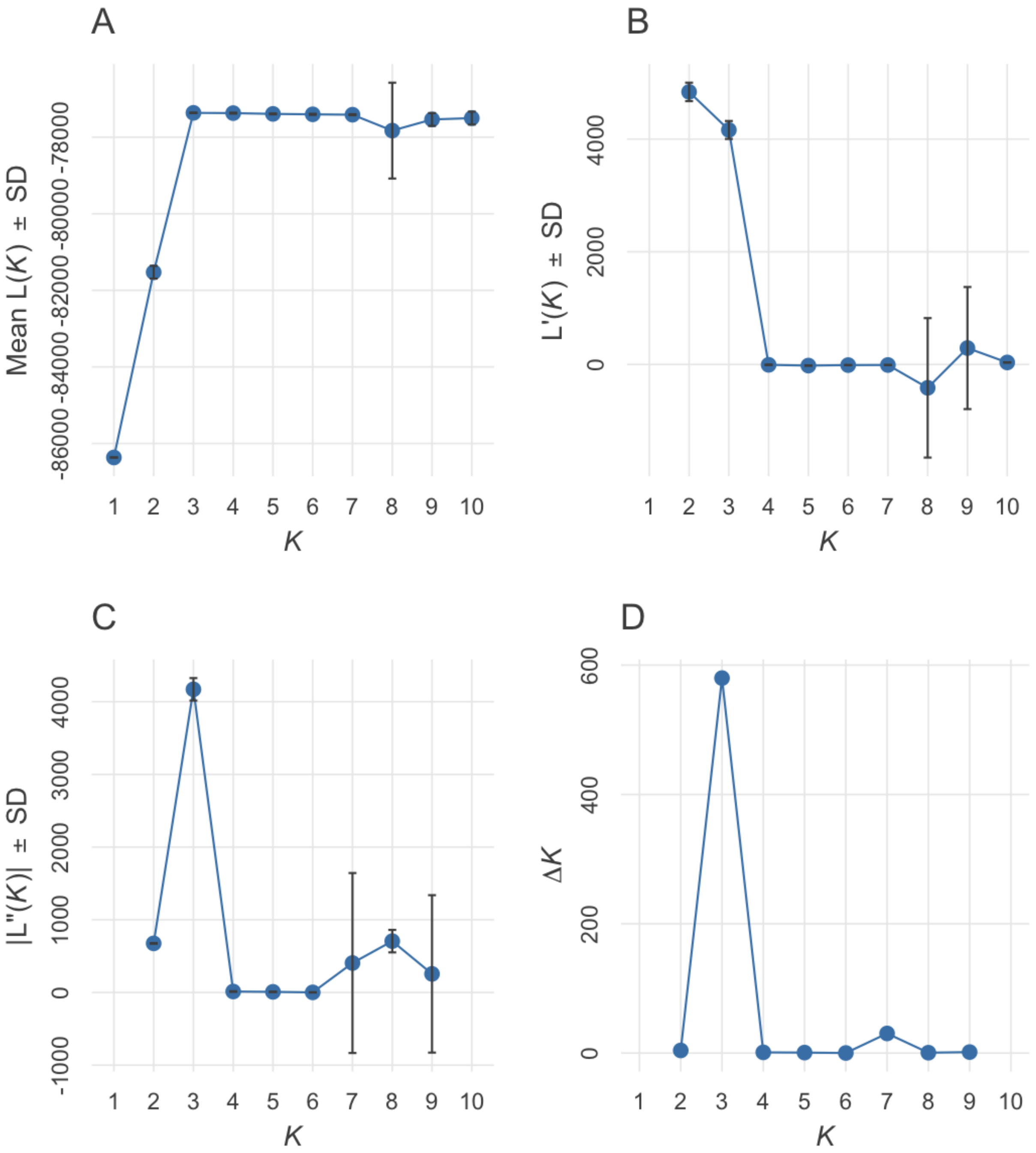

**Supplemental Figure S9.** Change in likelihood values from Structure results for the *D. elongata* SNP dataset, visualized according to four methods (A-D) for values of $K$ = 1 to $K$ = 10.

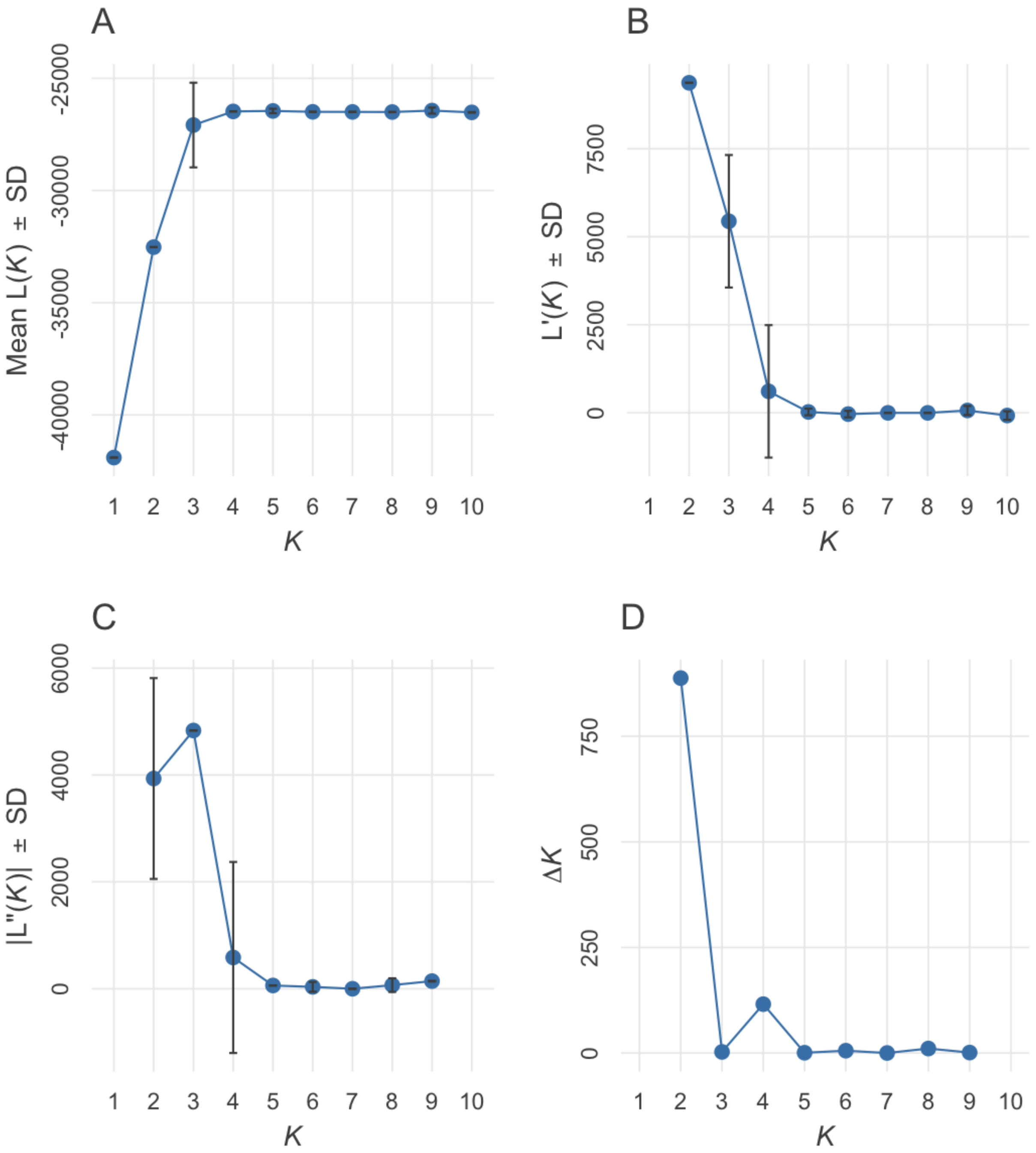

**Supplemental Figure S9.** Estimated ancestry according to latitude for individuals from sites indicated by color and shape of A) *D. sublineata* from the global set of Structure results and B) *D. carinulata* Fukang ecotype from the *D. carinulata* substructure results. The linear model for each distribution is plotted as a red line with standard error in grey shading.

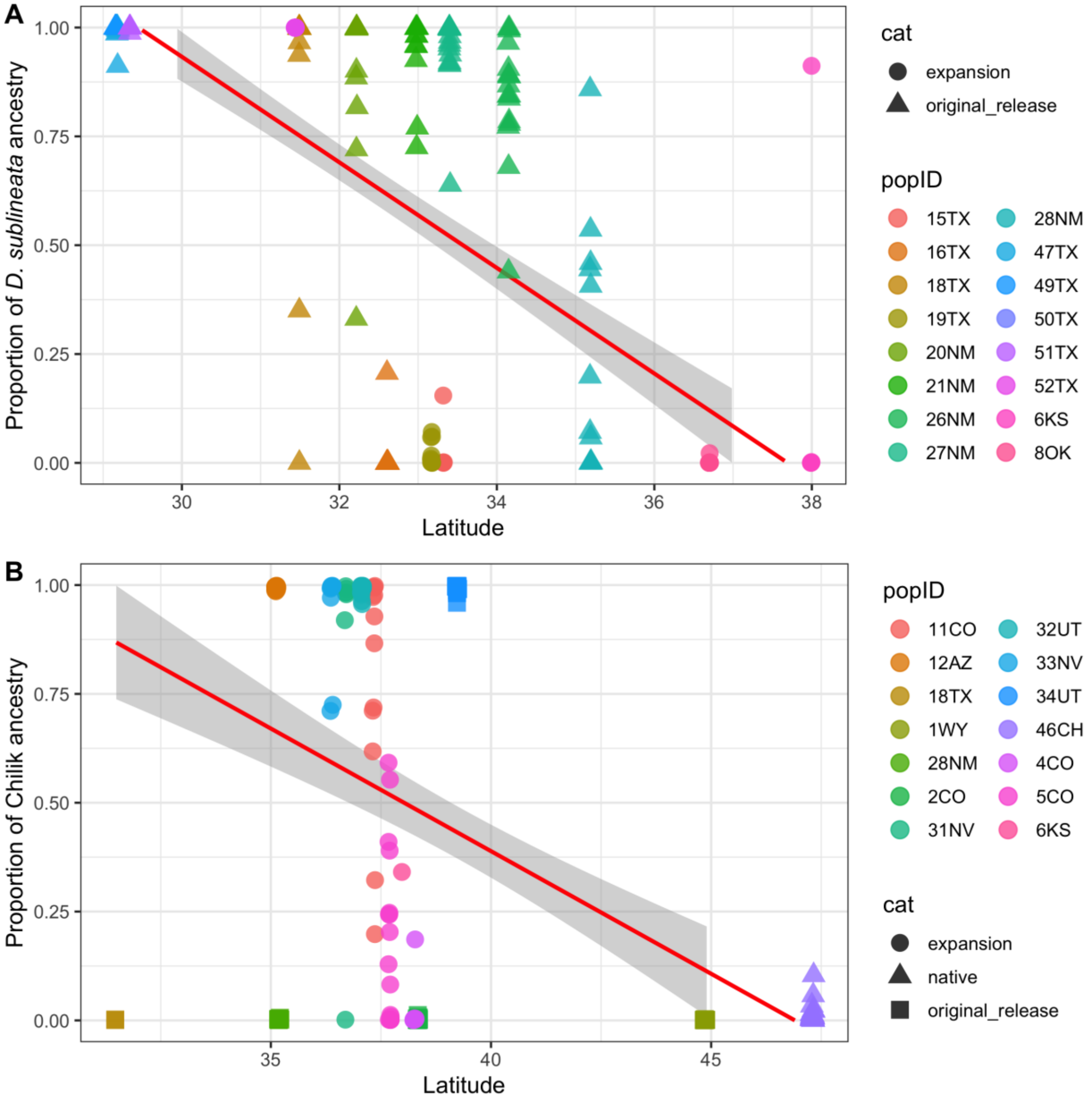

**Supplemental Figure S10.** Pairwise-$F_{ST}$ compared to haversine distance between respective sites indicate patterns of isolation-by-distance associated with population structure for *D. elongata* collected from the native range in Greece (Mantel's r = 0.086, $p$ = 0.492). Crossed-squares indicate pair-wise comparisons within ecotype, solid squares across ecotype. A linear model for each distribution projected behind points as a red line with standard error in grey to aid in visualization.

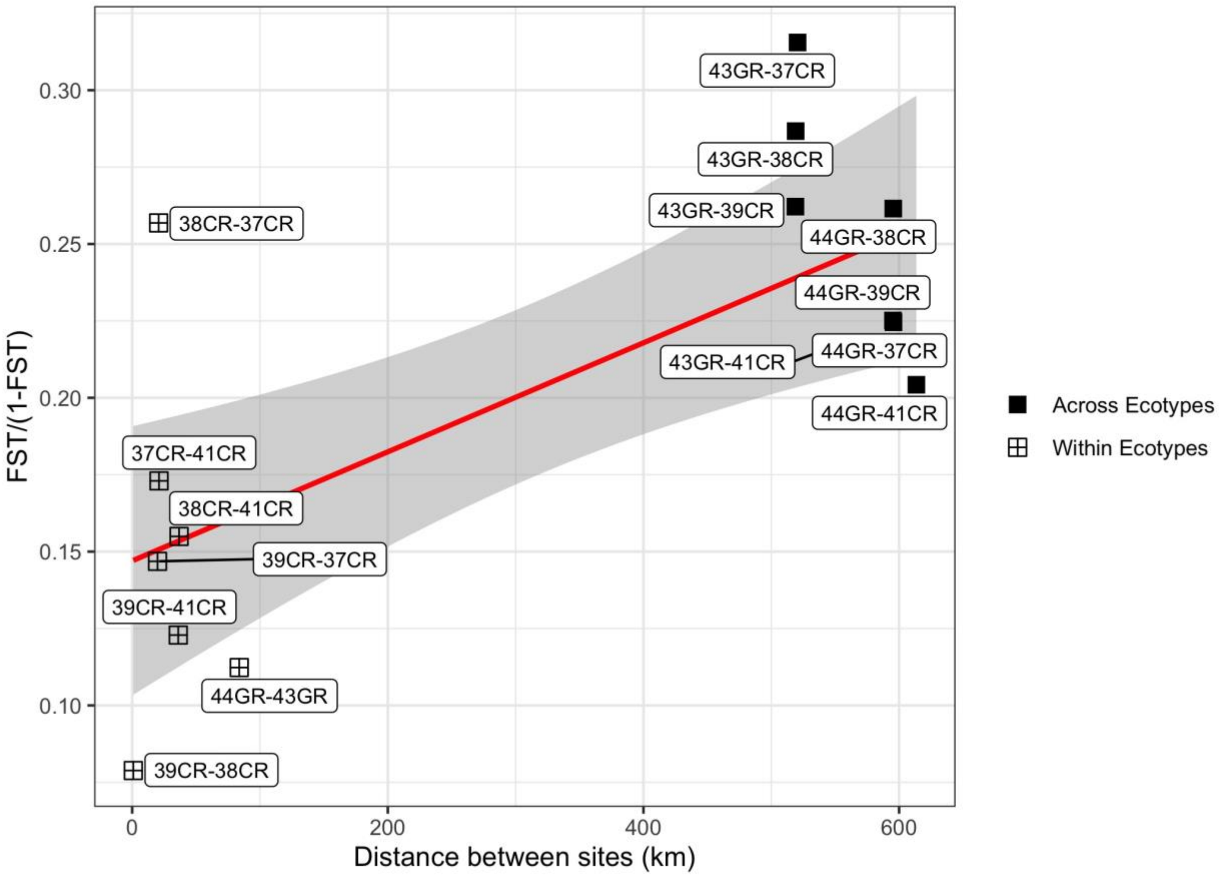

**Supplemental Figure S11.** Directionality index (ψ) for both *D. carinulata* ecotypes (origins) in North America. Positive stepwise increasing values suggest a directional founding effect. Each ecotype is outlined by a box, with admixture zones (Fig. 3) overlapping.

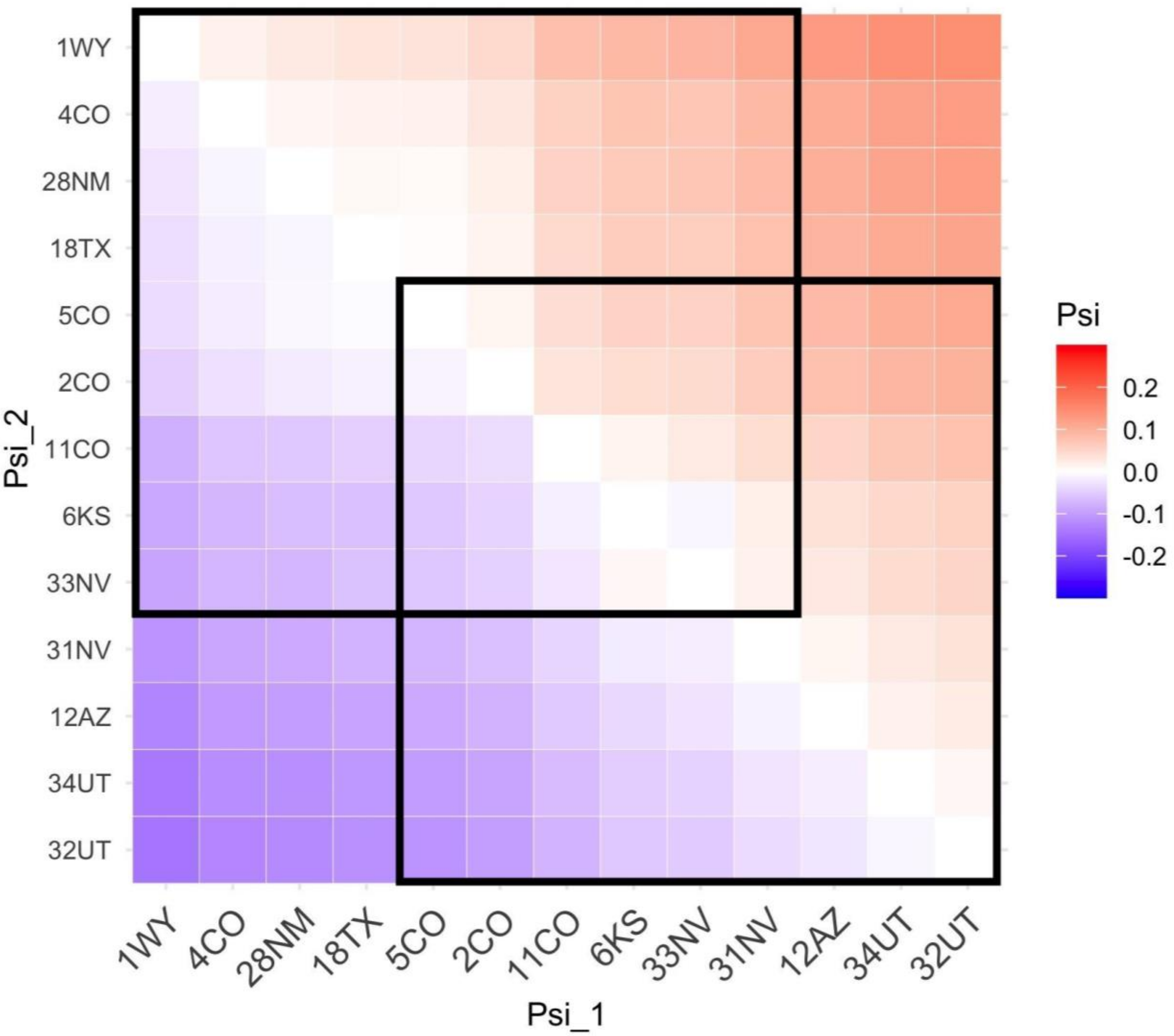
